# Tau pathology in cognitively normal older adults

**DOI:** 10.1101/611186

**Authors:** Jacob Ziontz, Murat Bilgel, Andrea T. Shafer, Abhay Moghekar, Wendy Elkins, Jessica Helphey, Gabriela Gomez, Danielle June, Michael A. McDonald, Robert F. Dannals, Babak Behnam Azad, Luigi Ferrucci, Dean F. Wong, Susan M. Resnick

**Affiliations:** Brain Aging and Behavior Section, Laboratory of Behavioral Neuroscience, National Institute on Aging, Baltimore, MD, USA; Longitudinal Studies Section, Translational Gerontology Branch, National Institute on Aging, Baltimore, MD, USA; Department of Radiology and Radiological Science, Johns Hopkins University School (JHU) of Medicine, Baltimore, MD, USA; Department of Psychiatry and Behavioral Sciences, JHU School of Medicine, Baltimore, MD, USA; Department of Neuroscience, JHU School of Medicine, Baltimore, MD, USA; Department of Neurology, JHU School of Medicine, Baltimore, MD, USA

**Keywords:** tau, AV-1451, T807, flortaucipir, FTP, PET, cognition, volume, longitudinal, cognitively normal

## Abstract

**INTRODUCTION:** Tau pathology, a hallmark of Alzheimer’s disease, is observed in the brains of virtually all individuals over 70.

**METHODS:** Using ^18^F-AV-1451 (^18^F-flortaucipir) PET, we evaluated tau pathology in 54 cognitively normal participants (mean age 77.5, SD 8.9) from the Baltimore Longitudinal Study of Aging. We assessed associations between PET signal and age, sex, race, and amyloid positivity. We investigated relationships between regional signal and retrospective rates of change in regional volumes and cognitive function adjusting for age, sex, and amyloid status.

**RESULTS:** Greater age, male sex, black race, and amyloid positivity were associated with higher ^18^F-AV-1451 retention in distinct brain regions. Retention in the entorhinal cortex was associated with lower entorhinal volume (*β* = −1.124, SE = 0.485, *p* = 0.025) and a steeper decline in memory performance (*β* = −0.086, SE = 0.039, *p* = 0.029).

**DISCUSSION:** Assessment of medial temporal tau pathology will provide insights into early structural brain changes associated with later cognitive impairment and Alzheimer’s disease.

## 1. Introduction

Pathological tau is a hallmark of several neurodegenerative diseases, most notably Alzheimer’s disease (AD). Tau promotes assembly and stability of microtubules in the nervous system [1], but its hyperphosphorylation leads to the formation of neurofibrillary tangles (NFT), which are observed at autopsy in brains of almost all 5 individuals above 70 [2]. NFTs in the medial temporal lobe in the absence of amyloid deposition and clinical symptomatology has been referred to as primary age-related tauopathy (PART) [3], though it is not clear if PART is distinct from the continuum of AD [4].

Tau pathology is hypothesized to be one of the earliest pathophysiological changes [5] in preclinical AD, spreading from the entorhinal cortex and hippocampus to the neocortex at later disease stages [6]. Positron emission tomography (PET) tau radiotracers have enabled the *in vivo* characterization of pathological tau. Cross-sectional studies including cognitively normal (CN) participants in addition to those with mild cognitive impairment (MCI) and AD indicate that lower cognitive performance is associated with higher tau tracer retention in the temporal lobe [7, 8] and the neocortex [9]. Lower hippocampal volume has also been associated with greater tau radiotracer retention in the hippocampus [8, 10]. Longitudinal studies including individuals ranging from CN to demented have further found that higher baseline tau tracer retention is associated with greater rates of brain volume loss [11, 12] and global cognitive decline [13–15].

Though pathological tau burden has been shown to be related to cognition and brain volume in individuals across the AD spectrum, the influence of tau in CN individuals who may or may not go on to develop clinical impairment remains unclear. There is a growing body of literature on the characterization of tau pathology using PET in this population. Several studies have observed associations between tau PET and brain volume in CN individuals. Lower gray matter volume intensity has been associated with greater tau PET retention in the medial temporal lobe [16], and studies utilizing retrospective longitudinal MRI data have further demonstrated that cortical thinning is related to tau PET particularly in medial and lateral temporal areas [17, 18]. However, the relationship between tau PET and cognitive performance is less clear. Results in CN individuals indicate that medial temporal lobe tau PET is associated with lower cross-sectional episodic memory and steeper retrospective decline in episodic memory adjusting for age, sex, and amyloid burden [18, 19]. Other studies have not found associations between regional tau PET and cognition [20], or have shown that tau PET interacts with amyloid to predict greater episodic memory decline only in amyloid+ individuals [21].

The lack of consensus in studies of CN individuals indicates a further need to analyze tau PET in this population and examine its association with brain volume loss and cognitive decline. The main goal of the current study was to study determine whether age- and amyloid-related differences in tau deposition can be detected among CN older adults, and if so, to investigate whether tau deposition is associated with regional brain volume and cognitive performance changes among CN. We first investigated factors associated with tau tracer retention in a sample of 54 CN older adults from the Baltimore Longitudinal Study of Aging (BLSA). We then examined regional tracer retention in regions where early tau accumulation is known to occur (i.e., the entorhinal cortex, hippocampus, and inferior temporal gyrus) in relation to retrospective longitudinal brain volume (over the course of about 7 years) and cognitive changes (over the course of about 13 years). We hypothesized that age and amyloid positivity would be associated with tau tracer retention given the relevance of these two factors for pathology spread, and that tau tracer retention would explain retrospective brain volume and cognitive decline in areas known to be affected early in AD. Understanding these relationships in a CN sample will allow us to better evaluate tau as a potential target for study and interventions to address brain aging and disease.

## 2. Methods

### 2.1. Participants

The study sample included CN BLSA participants with a ^18^F-AV-1451 (^18^F-flortaucipir) tau PET, a ^11^ C-Pittsburgh compound B (^11^C-PiB) amyloid PET within 2.2 years of tau PET, and a structural MRI. As of January 18, 2018, tau PET scans were acquired on 63 participants. Four had a non-CN status, two did not have an MRI at the time of analysis, one was subsequently discovered to have had an unreported myocardial infarction prior to enrollment (therefore meeting the exclusion criteria for PET study enrollment), and one was determined to be an outlier due to highly lateralized cortical signal. The final sample, after excluding these cases, consisted of 54 individuals (Table 1). For 47 of these participants, MRI and PET scans were ≤ 6 months apart. They were 0.6, 2.1, 2.1, 4.1, 4.6, 5.8, 7.3 years apart for the remaining 7 participants.

CN status was based on either (i) a Clinical Dementia Rating score [22] of zero and ≤ 3 errors on the Blessed Information-Memory-Concentration Test [23], and therefore the participant did not meet criteria for consensus conference; or (ii) the participant met criteria for consensus conference and was determined to be CN based on thorough review of clinical and neuropsychological data.

**Table 1:** Participant demographics.

| *a. Voxel-wise analysis sample (n = 54)* | **Amyloid−**<br>**(n = 41)** | **Amyloid+**<br>**(n = 13)** |
| --- | --- | --- |
| Age at <sup>18</sup>F-AV-1451 PET scan (yrs), mean (SD) | 77.2 (8.9) | 78.2 (9.3) |
| Male, n (%) | 16 (39%) | 8 (62%) |
| Black, n (%) | 8 (20%) | 4 (31%) |
| Yrs of education, mean (SD) | 17.9 (2.1) | 16.6 (2.2) |
| *APOE* ε4+, n (%) | 11 (27%) | 7 (54%) |
| *b. Cognition sample (n = 53)* | **Amyloid−**<br>**(n = 40)** | **Amyloid+**<br>**(n = 13)** |
| Age at <sup>18</sup>F-AV-1451 PET scan (yrs), mean (SD) | 77.6 (8.7) | 78.2 (9.3) |
| Male, n (%) | 16 (40%) | 8 (62%) |
| Black, n (%) | 8 (20%) | 4 (31%) |
| Yrs of education, mean (SD) | 17.8 (2.1) | 16.6 (2.2) |
| *APOE* ε4+, n (%) | 10 (25%) | 7 (54%) |
| Number of cognitive assessments, mean (SD) | 8 (5.2) | 6.5 (5.5) |
| Duration of cognitive follow-up (yrs), mean (SD) | 13.9 (8) | 11.5 (8.7) |
| *c. Brain volume sample (n = 50)* | **Amyloid−**<br>**(n = 38)** | **Amyloid+**<br>**(n = 12)** |
| Age at <sup>18</sup>F-AV-1451 PET scan (yrs), mean (SD) | 77.3 (8.9) | 77.8 (9.5) |
| Male, n (%) | 15 (39%) | 7 (58%) |
| Black, n (%) | 8 (21%) | 3 (25%) |
| Yrs of education, mean (SD) | 17.9 (2.1) | 16.7 (2.3) |
| *APOE* ε4+, n (%) | 10 (26%) | 6 (50%) |
| Number of MRI scans, mean (SD) | 5.3 (4.9) | 4.3 (4.6) |
| Duration of MRI follow-up (yrs), mean (SD) | 7.4 (6.8) | 6.2 (6.5) |

Research protocols were approved by local institutional review boards, and all participants gave written informed consent at each visit. At enrollment into the PET neuroimaging substudy of the BLSA, all participants were free of CNS disease (dementia, stroke, bipolar illness, epilepsy), severe cardiac disease, severe pulmonary disease, and metastatic cancer. One participant had a myocardial infarction and another was diagnosed with congestive heart failure after PET substudy enrollment but before tau PET scan.

### 2.2. Structural imaging

Magnetization-prepared rapid gradient echo (MPRAGE) images were acquired on a 3 T Philips Achieva scanner (repetition time = 6.8 ms, echo time = 3.2 ms, flip angle = 8^∘^, image matrix = 256 × 256, 170 slices, voxel size =1 × 1 × 1.2 mm). Anatomical labels and global and regional brain volumes were obtained using Multi-atlas region Segmentation using Ensembles of registration algorithms and parameters (MUSE) [24]. We performed intracranial volume (ICV) correction using the approach employed by Jack et al. [25], computing residual volumes for each region, which is the difference, in cm^3^, from the regional volume that would be expected at a given ICV.

### 2.3. PET imaging

Amyloid PET imaging is described in Appendix B. Tau PET scans were obtained over 30 min on a Siemens High Resolution Research Tomograph (HRRT) scanner starting 75 mins after an intravenous bolus injection of approximately 370 MBq (10 mCi) of ^18^F-AV-1451. Dynamic images were reconstructed using ordered subset expectation-maximization to yield 6 time frames of 5 mins each with approximately 2.5 mm full-width at half-maximum (FWHM) at the center of the field of view (image matrix = 256 × 256, 207 slices, voxel size = 1.22 × 1.22 × 1.22 mm). We aligned the time frames between 80–100 minutes to the first frame in this interval. The 20-min average PET image was registered onto the inhomogeneity-corrected MPRAGE using rigid registration. Anatomical labels defined in MRI space were transformed into PET space. The 20-min average PET image was partial volume corrected using the Region-Based Voxel-wise (RBV) method [26].

For the geometric transfer matrix step of RBV, we used 26 bilateral MUSE regions (see Appendix D). We computed standardized uptake value ratio (SUVR) images by dividing the partial volume corrected PET intensities by the mean within the inferior cerebellar gray matter, which was defined using the approach described by Baker et al. [27] based on the SUIT atlas [28]. We computed the average SUVR in three regions of interest (ROIs) corresponding to early stages of tau pathology: the entorhinal cortex, hippocampus, and inferior temporal gyrus (ITG). SUVR images were mapped into MNI space using the warp computed from deformable registration of the corresponding MRIs to a study-specific MRI template, and smoothed with a Gaussian filter (6 mm FWHM) prior to statistical analysis.

### 2.4. Neuropsychological testing

Cognitive domain scores were obtained for memory (California Verbal Learning Test (CVLT) [29] immediate and long-delay free recall), attention (Trail Making Test [30] Part A and Digit Span [31] Forward), executive function (Trail Making Test Part B and Digit Span Backward), fluency (Category [32] and Letter Fluency [33]), visuospatial processing (Card Rotations Test [34], Clock Drawing Test [35]), and processing speed (Digit Symbol Substitution Test) [31]. To obtain domain scores, each test score was first converted to a *z*-score using the baseline mean and standard deviation, and these *z*-scores were averaged within each cognitive domain. Prior to computing the *z*-scores for Trail Making Test Parts A and B, the individual cognitive test scores (time to completion, in seconds) were log transformed and negated so that higher *z*-scores indicated shorter time to completion.

### 2.5. Statistical analysis

#### 2.5.1. Factors associated with tau accumulation

We used multiple linear regression to assess the associations between demographics, amyloid positivity and voxel-wise ^18^F-AV-1451 SUVR. Independent variables included age, sex, race, amyloid status, and age × amyloid status. Education was not included as a predictor because of its low variance in our sample. Each independent variable was mean-centered to facilitate interpretation of model results. Voxel-wise linear regression was conducted using SPM12. Statistical significance was based on two-tailed T-tests with *p* < 0.001 (uncorrected for multiple comparisons) and restricted to clusters of ≥ 400 voxels.

#### 2.5.2. Longitudinal regional brain volume change and co-localized tau accumulation

Using separate linear mixed effects models for each of the three PET ROIs (entorhinal cortex, hippocampus, and ITG), we assessed the associations between regional ^18^F-AV-1451 SUVR and retrospective change in the volume of the same region (3 models total). The dependent variable was ICV-adjusted regional volumes prior to and concurrent with the tau PET scans. Age at and time from tau PET scan, sex, amyloid status (+ vs −), amyloid status × time, regional ^18^F-AV-1451 SUVR, and SUVR × time were included as independent variables. To facilitate interpretation, regional ^18^F-AV-1451 SUVRs were mean-centered. Random effects were included for intercept and time. This analysis was restricted to individuals who had a volumetric measurement prior to and within 3 years of tau PET (n = 50, total number of longitudinal MRI assessments = 253). Longitudinal MRI time points per participant relative to the time of tau PET scan are shown in Figure A.1. Statistical significance was based on two-tailed T-tests with *p* < 0.05 (uncorrected for multiple comparisons).

#### 2.5.3. Longitudinal cognition and tau accumulation

Using separate linear mixed effects models for each of the six cognitive domains and each of the three ROIs, we assessed associations between regional ^18^F-AV-1451 SUVR and retrospective change in cognition (18 models total). Age at and time from tau PET scan, sex, years of education, amyloid status, amyloid status × time interaction, regional ^18^F-AV-1451 SUVR, and regional SUVR × time interaction were included as independent variables. This analysis was restricted to individuals who had a cognitive assessment prior to and within 3 years of tau PET (n = 53). Longitudinal cognitive visits per participant relative to the time of tau PET scan are shown in Figure A.1. There were 401 total observations for memory, 351 for attention and executive function, 363 for fluency, 293 for visuospatial processing, and 249 for processing speed, with differences in sample sizes primarily reflecting historical differences in age at which specific tests were administered. Amyloid status and regional ^18^F-AV-1451 SUVRs were centered around the sample mean as before. Random effects were included for intercept and time. Statistical significance was based on two-tailed T-tests with *p* < 0.05 (uncorrected for multiple comparisons).

## 3. Results

### 3.1. Factors associated with tau accumulation

The association between age and ^18^F-AV-1451 SUVR was stronger among amyloid+ compared to amyloid–individuals in the right middle temporal gyrus, left middle frontal gyrus, and bilaterally in the cuneus, cingulate, superior frontal, and postcentral gyri (Table F.1). In the amyloid+ group, greater age was associated with higher ^18^F-AV-1451 SUVR in bilateral putamen, right inferior frontal, and right middle occipital gyri (Table F.2). Amyloid+ individuals had greater ^18^F-AV-1451 SUVR compared to amyloid-individuals in the right middle frontal gyrus, right superior and middle temporal gyri, left superior occipital gyrus, bilateral middle temporal gyri, middle occipital gyri, and cuneus (Table F.3). Men compared with women had higher ^18^F-AV-1451 SUVR in bilateral frontal, parietal, and lateral temporal cortices as well as in bilateral limbic areas (Table F.4). Black race was associated with higher ^18^F-AV-1451 SUVR in bilateral occipital and temporal lobes as well as superior frontal areas (Table F.5). These effects are visualized for select brain slices in Figure 1. Unthresholded statistical maps are available at https://neurovault.org/collections/LGNABWKB [36].

**Figure 1:**
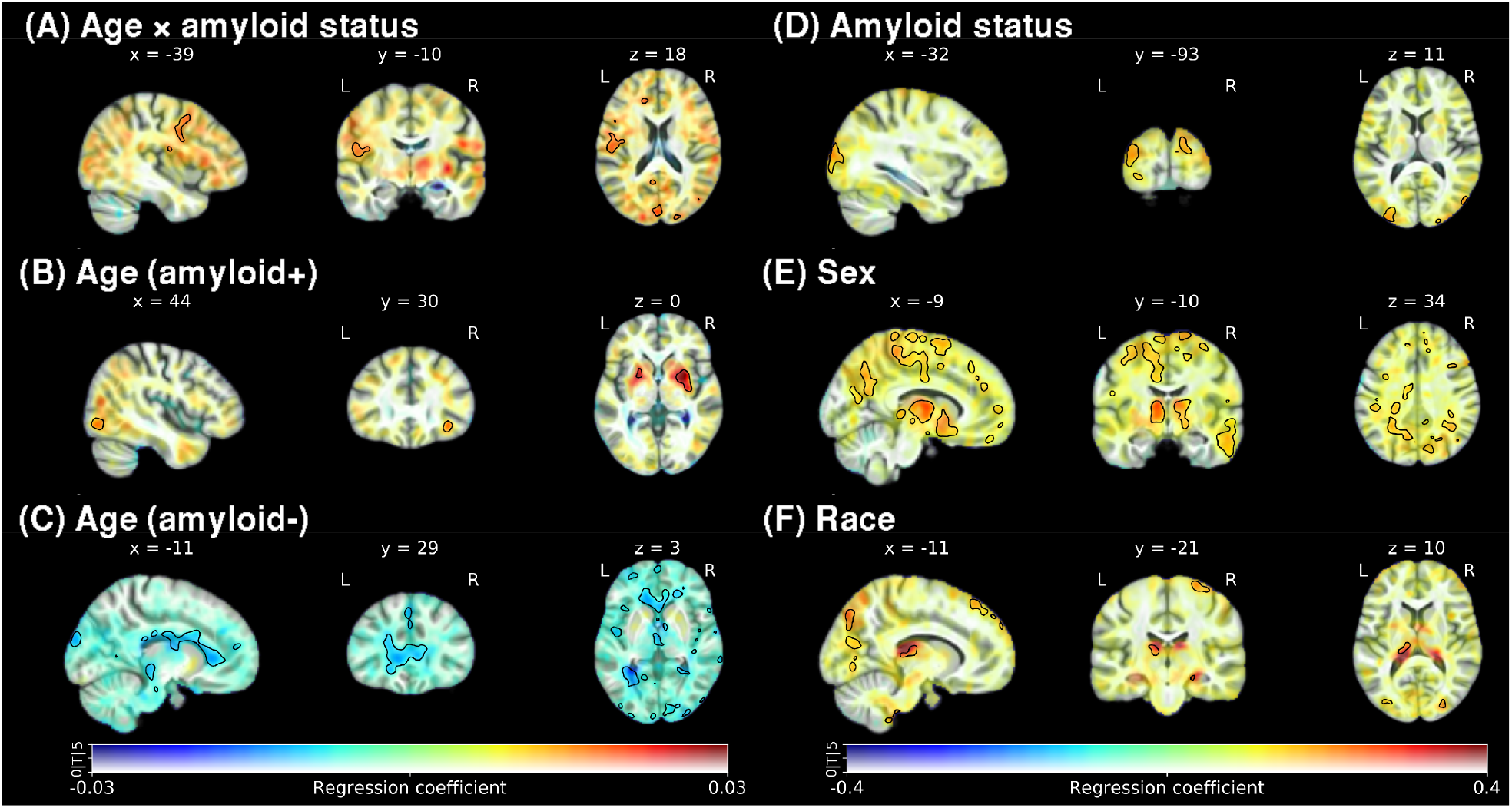
Predictors of ^18^F-AV-1451 tau tracer retention among cognitively normal older adults. In these dual-coded representations of voxel-wise linear regression results, color indicates the estimated regression coefficient (indicated along the horizontal axis of the colorbar) and transparency corresponds to the absolute T-value (with 0 as completely transparent and ≥ 5 as completely opaque, as indicated along the vertical axis of the colorbar). Voxels that reach significance (uncorrected *p* < 0.001, cluster size ≥ 400 voxels) are circumscribed by black contour to help with the interpretation of transparency. (A) Age by amyloid status interaction. (B) Main effect of age in amyloid+ individuals. (C) Main effect of age in amyloid– individuals. (D) Main effect of amyloid positivity. (E) Main effect of male sex. (F) Main effect of black race. Color bars on the left and right correspond to panels A–C and D–F, respectively.

### 3.2. Regional brain volume and tau accumulation

Cross-sectionally, higher entorhinal cortex ^18^F-AV-1451 SUVR was associated with smaller volume in this region (*β* = −1.124, SE = 0.485, *p* = 0.025) (Table 2). We did not find any associations between tau and volume in the hippocampus or ITG.

**Table 2:** Linear mixed effects models of the relationship between entorhinal ^18^F-AV-1451 SUVR and intracranial volume adjusted entorhinal cortex volume. Estimated fixed effects are reported along with their standard errors in parentheses.

| | *Dependent variable:*<br>Entorhinal volume ($\text{cm}^3$) |
| --- | --- |
| Intercept | $-0.301^{***}$ (0.063)<br>$p = 0.000004$ |
| Age at PET scan | $-0.030^{***}$ (0.007)<br>$p = 0.0002$ |
| Sex<br>(ref = female) | $0.338^*$ (0.129)<br>$p = 0.013$ |
| Amyloid group<br>(ref = amyloid-) | $0.231$ (0.152)<br>$p = 0.136$ |
| Entorhinal SUVR | $-1.124^*$ (0.485)<br>$p = 0.025$ |
| Time from PET | $-0.055^{***}$ (0.007)<br>$p = 0.000$ |
| Amyloid group<br>× time | $-0.001$ (0.017)<br>$p = 0.960$ |
| Entorhinal SUVR<br>× time | $-0.047$ (0.046)<br>$p = 0.308$ |
| *Note:* $^*p < 0.05$; $^{**}p < 0.01$; $^{***}p < 0.001$ | |

### 3.3. Cognition and tau accumulation

Cross-sectionally, greater ^18^F-AV-1451 SUVR in the hippocampus was associated with lower memory (*β* = −0.833, SE = 0.405, *p* = 0.046), and greater ^18^F-AV-1451 SUVR in the ITG was associated with lower attention scores (*β* = −2.451, SE = 1.046, *p* = 0.024) (Table 3).

**Table 3:**
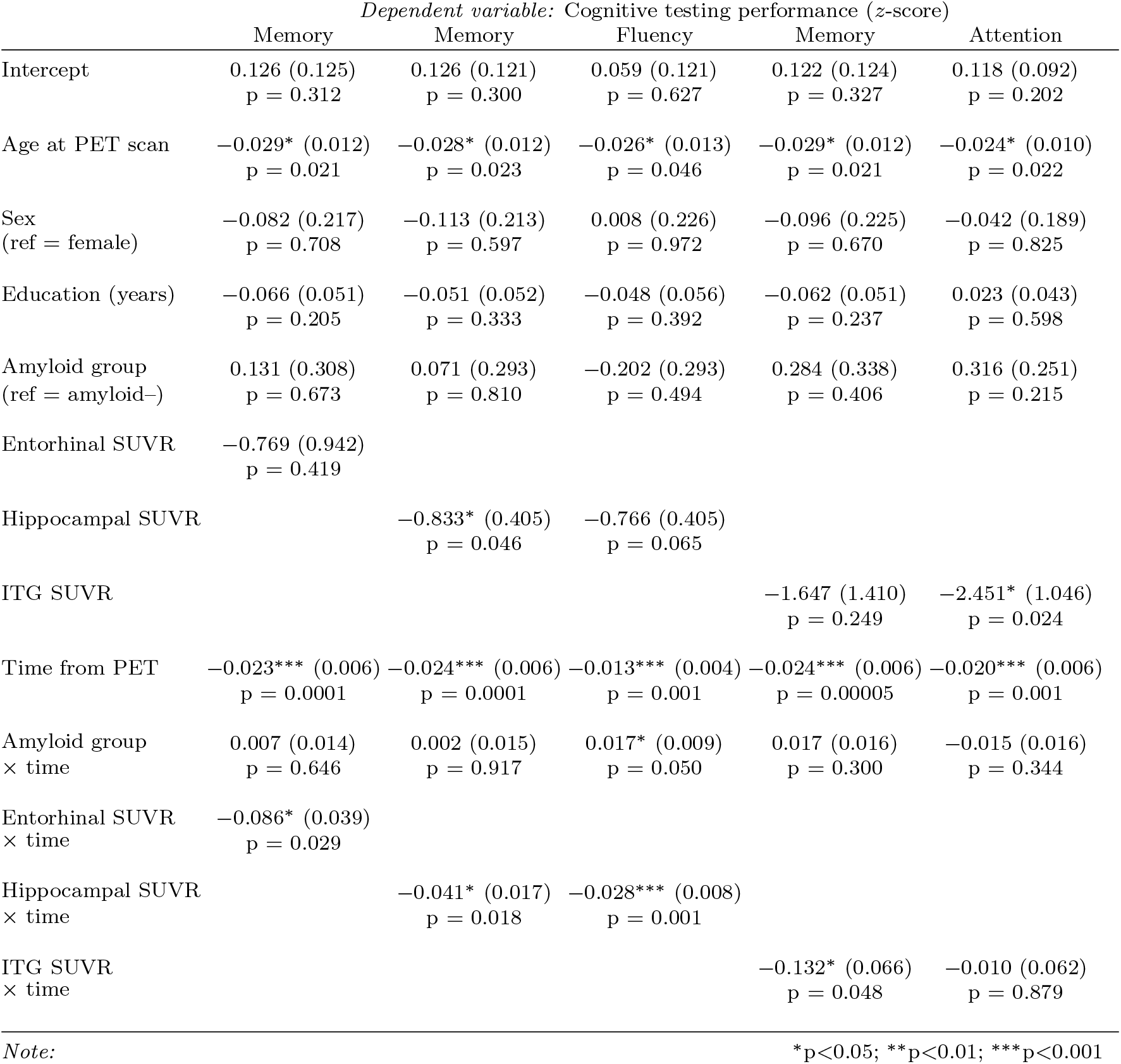
Linear mixed effects models of the relationship between entorhinal and hippocampal ^18^F-AV-1451 SUVR and cognition. Each column represents a separate model. Each model includes tau measured in a single ROI. Estimated fixed effects are reported along with their standard errors in parentheses.

|  | *Dependent variable: Cognitive testing performance (z-score)* |  |  |  |  |
| --- | --- | --- | --- | --- | --- |
|  | Memory | Memory | Fluency | Memory | Attention |
| Intercept | 0.126 (0.125)<br>p = 0.312 | 0.126 (0.121)<br>p = 0.300 | 0.059 (0.121)<br>p = 0.627 | 0.122 (0.124)<br>p = 0.327 | 0.118 (0.092)<br>p = 0.202 |
| Age at PET scan | -0.029* (0.012)<br>p = 0.021 | -0.028* (0.012)<br>p = 0.023 | -0.026* (0.013)<br>p = 0.046 | -0.029* (0.012)<br>p = 0.021 | -0.024* (0.010)<br>p = 0.022 |
| Sex<br>(ref = female) | -0.082 (0.217)<br>p = 0.708 | -0.113 (0.213)<br>p = 0.597 | 0.008 (0.226)<br>p = 0.972 | -0.096 (0.225)<br>p = 0.670 | -0.042 (0.189)<br>p = 0.825 |
| Education (years) | -0.066 (0.051)<br>p = 0.205 | -0.051 (0.052)<br>p = 0.333 | -0.048 (0.056)<br>p = 0.392 | -0.062 (0.051)<br>p = 0.237 | 0.023 (0.043)<br>p = 0.598 |
| Amyloid group<br>(ref = amyloid-) | 0.131 (0.308)<br>p = 0.673 | 0.071 (0.293)<br>p = 0.810 | -0.202 (0.293)<br>p = 0.494 | 0.284 (0.338)<br>p = 0.406 | 0.316 (0.251)<br>p = 0.215 |
| Entorhinal SUVR | -0.769 (0.942)<br>p = 0.419 |  |  |  |  |
| Hippocampal SUVR |  | -0.833* (0.405)<br>p = 0.046 | -0.766 (0.405)<br>p = 0.065 |  |  |
| ITG SUVR |  |  |  | -1.647 (1.410)<br>p = 0.249 | -2.451* (1.046)<br>p = 0.024 |
| Time from PET | -0.023*** (0.006)<br>p = 0.0001 | -0.024*** (0.006)<br>p = 0.0001 | -0.013*** (0.004)<br>p = 0.001 | -0.024*** (0.006)<br>p = 0.00005 | -0.020*** (0.006)<br>p = 0.001 |
| Amyloid group<br>× time | 0.007 (0.014)<br>p = 0.646 | 0.002 (0.015)<br>p = 0.917 | 0.017* (0.009)<br>p = 0.050 | 0.017 (0.016)<br>p = 0.300 | -0.015 (0.016)<br>p = 0.344 |
| Entorhinal SUVR<br>× time | -0.086* (0.039)<br>p = 0.029 |  |  |  |  |
| Hippocampal SUVR<br>× time |  | -0.041* (0.017)<br>p = 0.018 | -0.028*** (0.008)<br>p = 0.001 |  |  |
| ITG SUVR<br>× time |  |  |  | -0.132* (0.066)<br>p = 0.048 | -0.010 (0.062)<br>p = 0.879 |
| *Note:* |  |  |  | *p<0.05; **p<0.01; ***p<0.001 |  |

Steeper decline in memory was associated with greater ^18^F-AV-1451 SUVR in the entorhinal cortex (*β* = −0.086, SE = 0.039, *p* = 0.029) (Figure 2), hippocampus (*β* = −0.041, SE = 0.017, *p* = 0.018), and ITG (*β* = −0.132, SE = 0.066, *p* = 0.048). (Table 3). The strength of this association was greater for the CVLT long-delay free recall component than for the immediate recall component for all three regions (Table G.4). In addition, steeper decline in fluency was associated with ^18^F-AV-1451 SUVR in the hippocampus (*β* = −0.028, SE = 0.008, *p* < 0.001).

**Figure 2:**
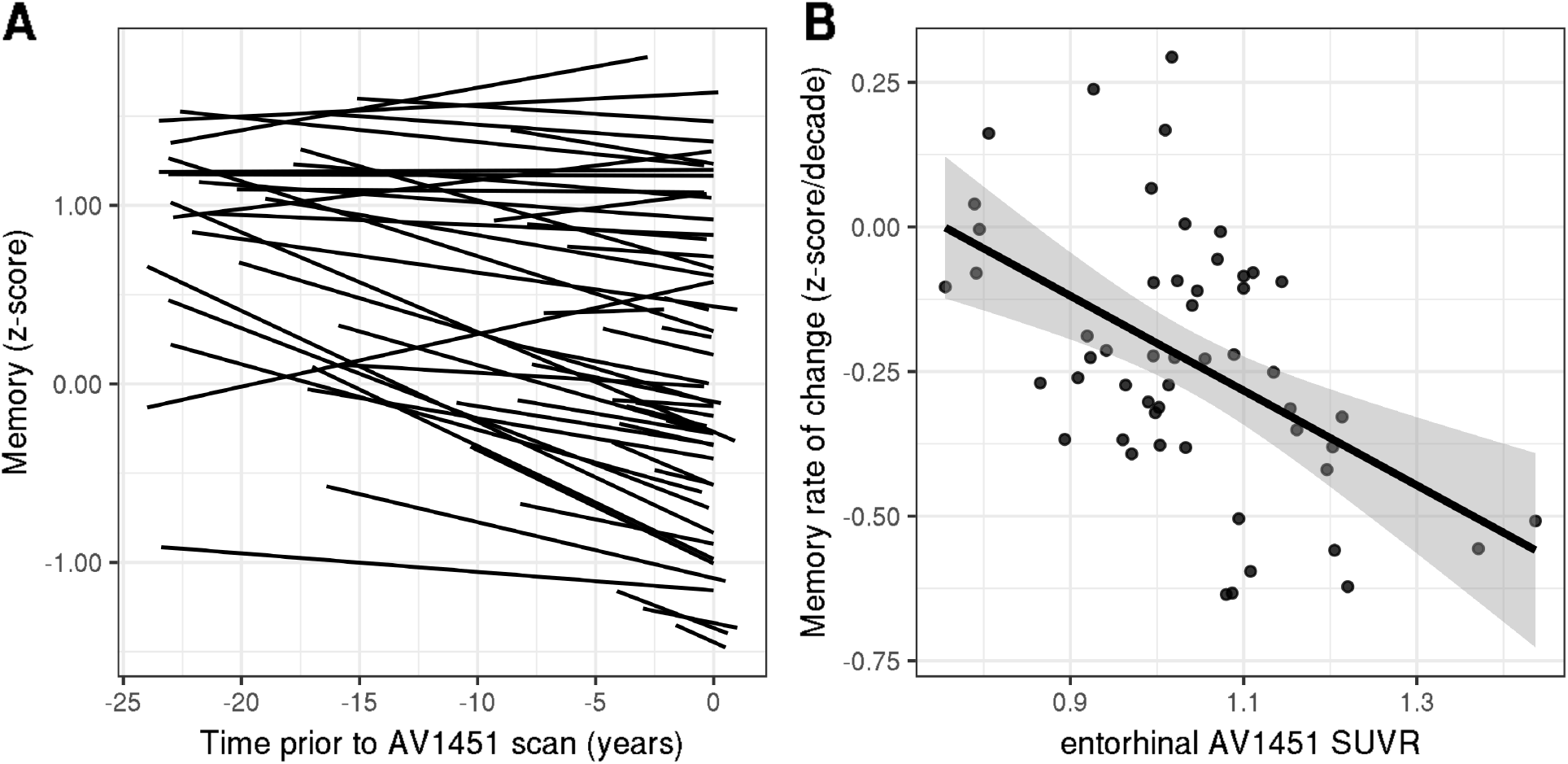
Entorhinal ^18^F-AV-1451 tau tracer retention is associated with steeper retrospective longitudinal decline in the composite memory score. (A) Individual-level memory change predicted by linear mixed effects model. (B) Rate of decline (z-score/decade) in memory performance as a function of ^18^F-AV-1451 tau tracer retention in the entorhinal cortex. Fitted values for rate of change are plotted for each individual in the sample.

The full set of results for each ROI and each cognitive domain are reported in Tables G.1, G.2, and G.3.

## 4. Discussion

Our study investigated tau tracer retention among cognitively normal individuals. We evaluated the cross-sectional associations of ^18^F-AV-1451 SUVR with age, sex, race, and amyloid status, and futher assessed the associations between ^18^F-AV-1451 SUVR and rates of volumetric and cognitive change. We found that higher ^18^F-AV-1451 retention in the entorhinal cortex was associated with lower volume in this region. We also found that greater ^18^F-AV-1451 retention in the entorhinal cortex, hippocampus, and inferior temporal gyrus were each associated with steeper decline in verbal memory.

Our finding of higher ^18^F-AV-1451 SUVR in temporal, temporoparietal, and frontal cortical areas among amyloid+ compared to amyloid–individuals is in agreement with previous studies of cognitively normal older adults [19, 37]. In the amyloid— group, ^18^F-AV-1451 SUVR was lower at greater ages in periventricular white matter and CSF, which might be due to age-related differences in radiotracer clearance among amyloid–individuals. Conversely, ^18^F-AV-1451 SUVR was higher at older ages among amyloid+ individuals in the putamen, right inferior frontal, and right middle occipital gyri. The association between tracer retention and age modulated by amyloid status did not reach significance in the putamen, but amyloid+ individuals exhibited stronger associations in several cortical regions. These findings suggest that the association in the putamen may be driven by non-specific binding whereas cortical associations may more likely be due to tau pathology. Similarly, the observed interaction between amyloid status and age might be reflective of cortical areas of faster tau accumulation among amyloid+ individuals. This interpretation is supported by the finding of a previous longitudinal tau PET study showing that amyloid+ individuals had steeper tau tracer retention increases in basal and mid-temporal, retrosplenial, posterior cingulate, and entorhinal cortex [38]. Another study of longitudinal tau accumulation showed increases in ^18^F-AV-1451 retention over 1—3 years in temporal and medial parietal areas in healthy older adults [39]. These findings were further expanded upon by a recent study reporting that individuals with baseline ^18^F-AV-1451 SUVRs in the second quartile exhibited tau tracer retention increases in inferior and lateral temporal cortex and in posterior cingulate over 18 months [15].

Men in our sample had higher ^18^F-AV-1451 SUVR than women, mainly in frontal and parietal white matter and thalamus. Previous studies utilizing tau PET imaging have not shown widespread or consistent sex differences in tracer retention [37, 40], and given that women exhibit a greater degree of AD pathology than men in *ex vivo* measures of tau [41], it seems likely that the sex differences we observed are largely driven by non-specific binding. Additionally, we found higher ^18^F-AV-1451 SUVR among black individuals in confined regions of the cortex. Black individuals exhibit lower levels of CSF-tau than white individuals [42], but greater incidence of postmortem NFT in Braak V/VI in black individuals has also been observed [43]. Race-related differences in ^18^F-AV-1451 retention in the choroid plexus have been previously reported [44], but the proximity of most statistically significant clusters to the edge of the brain in our sample suggests that these findings may be in part due to spill-over from non-specific binding of the tracer to meningeal neuromelanin. Potential sex and race differences in tau deposition will require further study in large and diverse samples.

Adjusting for age, sex, and amyloid status, we found that higher entorhinal ^18^F-AV-1451 SUVR was associated with lower brain volume in the entorhinal cortex. These results are in line with previous findings suggesting that tau accumulation may help explain differences in regional brain volumes while individuals are still cognitively normal [16–18], though these studies report more extensive associations.

Adjusting for age, sex, education, and amyloid status, we observed that greater ^18^F-AV-1451 retention in the entorhinal cortex, hippocampus, and inferior temporal gyrus was associated with steeper decline in verbal memory performance. This finding reinforces the notion that pathological tau in areas of early accumulation may influence changes in cognitive domains known to be affected in AD even in cognitively normal individuals. In a previous analysis using the BLSA amyloid PET data, we had reported an association between amyloid status and steeper memory decline [45]. Interestingly, this association was not statistically significant in our current analyses including regional ^18^F-AV-1451 SUVR as an independent variable. This suggests that tau rather than amyloid deposition may be more strongly associated with cognition, consistent with previous findings [7]. ITG tau PET retention was also associated with lower cross-sectional attention, and retention in the hippocampus was associated with steeper declines in fluency. Some studies have reported a relationship between tau burden and performance in cognitive domains other than memory [13, 46], though these findings may have been driven by the inclusion of clinically-impaired individuals, as this relationship is not consistently seen in CN individuals [21].

This study has several limitations. While the ^18^F-AV-1451 tracer has been demonstrated to have good specificity for tau tangles, it is also known to exhibit non-specific binding in the basal ganglia, choroid plexus, and to MAO-A, neuromelanin, and pigmented or mineralized vascular structures [47]. We did not perform multiple comparison correction given that a consensus method for correcting for dependent tests in longitudinal data sets is not available. We assessed the relationships between tau tracer retention and cognitive and volume declines retrospectively rather than prospectively due to the relatively recent implementation of tau PET in the BLSA. For this same reason, our sample size was limited, particularly for amyloid+ individuals. Future studies in a larger sample will be necessary to more fully investigate associations between amyloid, ^18^F-AV-1451 retention, and time.

Our study also has several strengths. The considerable amount of longitudinal cognitive testing and structural MRI data from the BLSA allowed us to assess the association between tau tracer retention and retrospective longitudinal decline in these measures over longer periods of time compared to other studies, and thereby yielding greater confidence in our estimates of rates of change. The spatial resolution of our tau PET scans, at approximately 2.5 mm FWHM at the center of the field of view, compares favorably to that of other studies of tau PET imaging among CN individuals. Better spatial resolution translates to less pronounced the partial volume effects, which is an advantage for image quantification, especially in medial temporal regions that are susceptible to spill-over from the choroid plexus.

Overall, our results point to a relationship between tau pathology and early changes in cognition in older individuals, even for those without a high degree of pathology or cognitive impairment. These findings also suggest the importance of ^18^F-AV-1451 PET for characterizing tau pathology in cognitively intact individuals and as a potential tool for predicting cognitive change early in AD progression. Future studies should investigate prospective cognitive and volumetric changes in relation to both timing and spread of tau deposition and their utility in predicting the trajectory of AD pathologies and symptoms. Effects of tau deposition on changes in other measures of brain integrity, such as brain networks underlying cognitive function, may provide additional insights into the relationship between pathology and cognitive decline in cognitively normal individuals.

## Abbreviations

AD: Alzheimer’s disease;
*APOE*: apolipoprotein E;
BA: Brodmann area;
BLSA: Baltimore Longitudinal Study of Aging;
CN: cognitively normal;
CSF: cerebrospinal fluid;
CVLT: California Verbal Learning Test;
DVR: distribution volume ratio;
FWHM: full-width at half-maximum;
HRRT: High Resolution Research Tomograph;
ICV: intracranial volume;
ITG: inferior temporal gyrus;
MCI: mild cognitive impairment;
MPRAGE: magnetization-prepared rapid gradient echo;
MRI: magnetic resonance imaging;
MUSE: multi-atlas region segmentation using ensembles of registration algorithms and parameters;
NFT: neurofibrillary tangle;
PART: primary age-related tauopathy;
PET: positron emission tomography;
PiB: Pittsburgh compound B;
RBV: region-based voxel-wise partial volume correction;
ROI: region of interest;
SD: standard deviation;
SE: standard error;
SUVR: standardized uptake value ratio

## 5. Acknowledgments

We thank the Baltimore Longitudinal Study of Aging participants and staff; the Laboratory of Behavioral Neuroscience Neuropsychology Testing Group; Noble George, Daniel Holt, Hong Fan, and the rest of the Johns Hopkins PET facility staff for their dedication to the BLSA studies and their assistance, and the Center for Biomedical Image Computing and Analytics for providing MUSE labels and their contributions to MRI analysis. We received valuable feedback from our colleagues at the National Institute on Aging, in particular from Dr. Lori Beason-Held on neuroanatomy, for which we are grateful. We also thank Avid Radiopharmaceuticals for enabling the use of the ^18^F-AV-1451 tracer and providing the precursor. Avid Radiopharmaceuticals did not provide direct funding for any of the PET studes nor personnel and were not involved in data analysis or interpretation.

## Appendix A. Longitudinal MRI volume and cognitive data included in linear mixed effects models

**Figure A.1:**
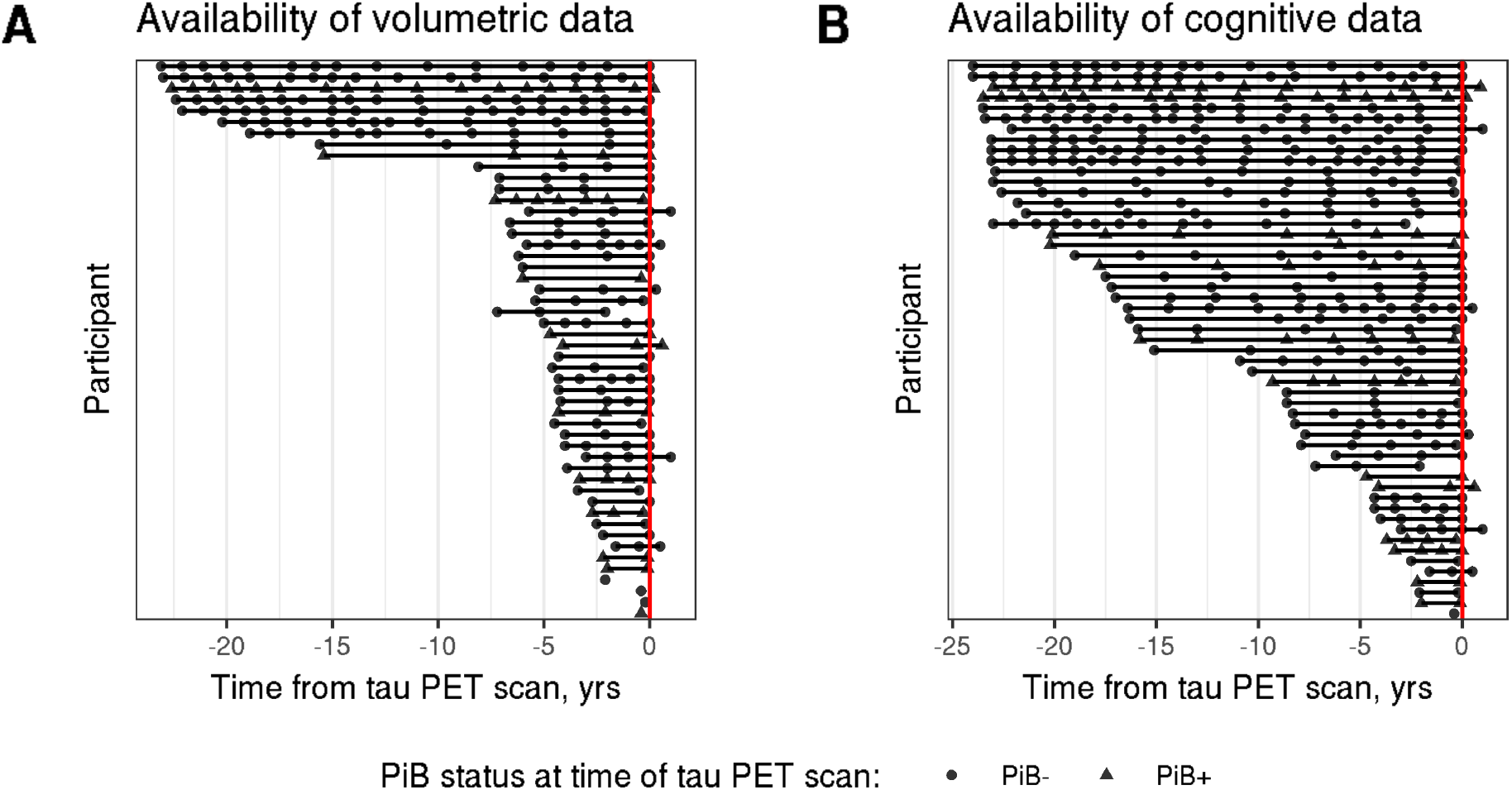
Availability of longitudinal A) volumetric and B) cognitive data per participant relative to the time of tau PET scan. Cognitive data availability differed across neuropsychological tests. Panel B shows the available data points for the CVLT. The red vertical line indicates the time of tau PET scan. Participants are sorted by their total follow-up duration (i.e., time between last and first volumetric or cognitive assessment).

## Appendix B. Amyloid PET imaging

Amyloid PET scans were obtained over 70 min on a GE Advance scanner immediately following an intravenous bolus injection of approximately 555 MBq (15 mCi) of ^11^-C-PiB. Dynamic images were reconstructed using filtered back-projection with a ramp filter, yielding a spatial resolution of approximately 4.5 mm FWHM at the center of the field of view (image matrix = 128 × 128, 35 slices, voxel size = 2 × 2 × 4.25 mm). Each of the 33 time frames was aligned to the mean of the first 2 min to correct for motion using SPM’s Realign (https://www.fil.ion.ucl.ac.uk/spm/software/spm12/) [48]. The average of the first 20 min of PET scans was rigidly registered onto the corresponding inhomogeneity-corrected MPRAGE, and the anatomical label image was transformed from MRI to PET space using FLIRT [49] implemented in FSL (https://fsl.fmrib.ox.ac.uk/fsl, version 6.0) [50]. Distribution volume ratio (DVR) images were computed in PET native space using a simplified reference tissue model [51] with cerebellar gray matter as the reference region. Mean cortical amyloid-*β* burden was calculated as the average of the DVR values in cingulate, frontal, parietal (including precuneus), lateral temporal, and lateral occipital cortical regions, excluding the sensorimotor strip. Individuals were categorized as amyloid −/+ based on a mean cortical DVR threshold of 1.057, which was derived from a Gaussian mixture model (Figure B.1).

**Figure B.1:**
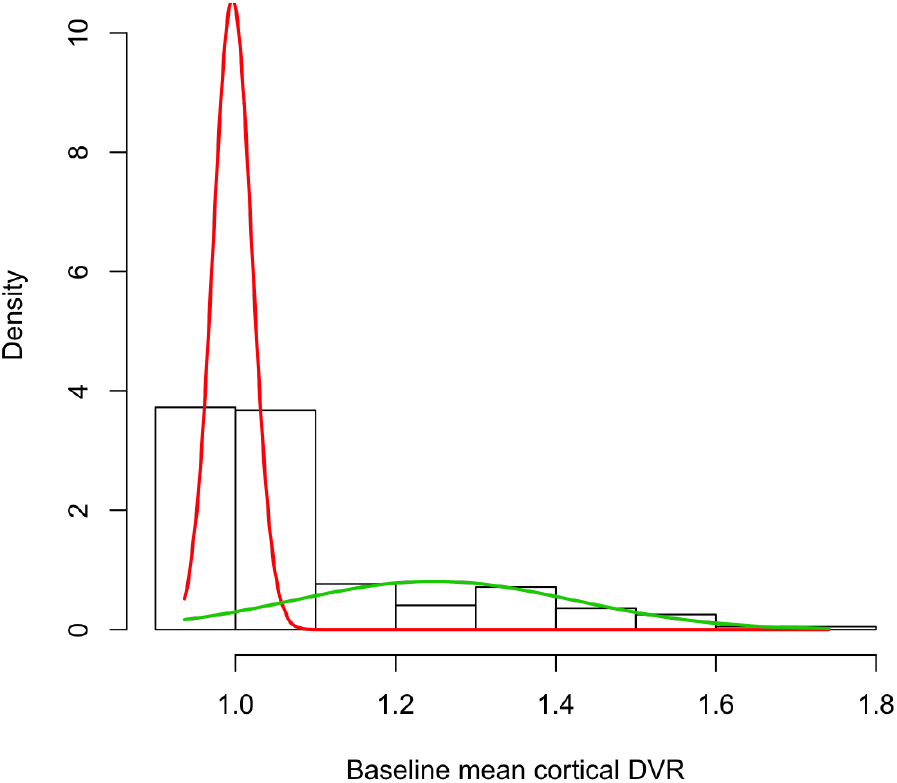
Result of the two-class Gaussian mixture model fitted on baseline mean cortical DVR. Red and green density curves correspond to the amyloid– and amyloid+ groups, respectively. Mean cortical DVR corresponding to the intersection of the two densities is the threshold for determining amyloid—/+ status.

## C. Software and computational reproducibility

### C.1. Software used for PET image processing

Time frame alignment of PET images was performed using SPM12’s Realign (https://www.fil.ion.ucl.ac.uk/spm/software/spm12/) [48]. Coregistration of PET and MRI images was performed with FLIRT [49] implemented in FSL (https://fsl.fmrib.ox.ac.uk/fsl, version 6.0) [50]. For partial volume correction, we used the PETPVC toolbox (https://github.com/UCL/PETPVC, version 1.2.0-b) [52]. For deformable registration of participant MRI images to the study-specific template, we used ANTs (http://stnava.github.io/ANTs/, version 2.1.0) [53]. PET image processing steps were streamlined using nipype (https://nipype.readthedocs.io/) [54] in Python 3.7.2.

### Appendix C.2. Software used for image visualization

Voxel-wise linear regression results were visualized as dual-coded images [55] using `nanslice` (https://github.com/spinicist/nanslice).

### Appendix C.3. Software used for statistical analyses

We used the `nlme` [56] package in R (https://cran.r-project.org, version 3.5.1) to fit the linear mixed effects models.

### Appendix C.4. Software used for writing the manuscript

To compile the manuscript, we used the following R packages: `knitr` [57, 58] to generate the manuscript directly incorporating results from R, `kableExtra` [59] to format tables, `stargazer` [60] to tabulate model results, `ggplot2` [61] to generate the scatter and trajectory plots, and `ggpubr` [62] to create panel figures.

### Appendix C.5. Computational reproducibility

PET image processing steps to generate SUVR images and all statistical analyses were containerized using Singularity [63] to ensure computational reproducibility. Code for replicating the statistical analyses and producing this manuscript is provided at https://gitlab.com/bilgelm/tau_predictors, and the Singularity image containing all necessary software at https://www.singularity-hub.org/collections/2612. Data used in these analyses are available upon request from the BLSA website (https://www.blsa.nih.gov). All requests are reviewed by the BLSA Data Sharing Proposal Review Committee and are also subject to approval from the NIH IRB.

## Appendix D. Regions used in partial volume correction of tau PET

Regions used for the geometric transfer matrix step of the RBV partial volume correction method applied to tau PET images were: background, ventricles and cerebrospinal fluid, basal ganglia, thalamus, brainstem, hippocampus, amygdala, cerebral white matter, inferior frontal gray matter, lateral frontal gray matter, medial frontal gray matter, opercular frontal gray matter, lateral parietal gray matter, medial parietal gray matter, fusiform, lateral temporal gray matter, supratemporal gray matter, inferior occipital gray matter, lateral occipital gray matter, medial occipital gray matter, limbic medial temporal gray matter, cingulate gray matter, insula gray matter, cerebellar white matter, cerebellar gray matter, cerebellar vermis.

## Appendix E. Regional analyses of predictors of ^18^F-AV-1451 SUVR

Given that voxel-wise analyses might be susceptible to inter-subject registration errors, we tested the relationship between these predictors and regional ^18^F-AV-1451 SUVR means computed in native PET space to verify our voxel-wise findings. Four anatomical regions were selected based on the observed voxel-wise effects and these effects were corroborated using a native space region of interest approach. To corroborate the observed voxel-wise effects of predictors of ^18^F-AV-1451 SUVR, we used linear regression to test the relationship between demographics, amyloid positivity, and regional ^18^F-AV-1451 SUVR. Regions within areas indicated by the voxel-wise analyses were selected to be tested for each predictor in the model. In agreement with voxel-wise results, amyloid status modulated the association of greater age with ^18^F-AV-1451 SUVR in the medial occipital gray matter (*β* = 0.009, SE = 0.003, *p* = 0.003). Amyloid positive individuals also exhibited greater ^18^F-AV-1451 SUVR in the bilateral inferior temporal gyrus compared to amyloid negative individuals (*β* = 0.095, SE = 0.028, *p* = 0.002). In addition, male sex was associated with greater ^18^F-AV-1451 SUVR in the frontal gray matter (*β* = 0.102, SE = 0.024, *p* < 0.001). Finally, black individuals showed greater ^18^F-AV-1451 retention in the superior frontal gyrus (*β* = 0.137, SE = 0.032, *p* < 0.001) (Table E.1).

**Table E.1:** Associations between participant demographics, amyloid status, and regional ^18^F-AV-1451 SUVR. Estimated fixed effects are reported along with their standard errors in parentheses.

|  | *Dependent variable: Regional $^{18}\text{F}$-AV-1451 SUVR* |  |  |  |
| --- | --- | --- | --- | --- |
|  | Medial occipital GM | Inferior temporal gyrus | Frontal gray matter | Superior frontal gyrus |
| Intercept | 1.139*** (0.011)<br>p = 0.000 | 1.243*** (0.012)<br>p = 0.000 | 1.199*** (0.011)<br>p = 0.000 | 1.165*** (0.013)<br>p = 0.000 |
| Age at PET scan | -0.002 (0.001)<br>p = 0.148 | 0.00002 (0.001)<br>p = 0.987 | -0.001 (0.001)<br>p = 0.530 | 0.001 (0.002)<br>p = 0.531 |
| Sex<br>(ref = female) | 0.046* (0.022)<br>p = 0.046 | 0.049* (0.024)<br>p = 0.050 | 0.102*** (0.024)<br>p = 0.0001 | 0.099*** (0.028)<br>p = 0.001 |
| Race<br>(ref = non-black) | 0.102*** (0.026)<br>p = 0.0004 | 0.065* (0.029)<br>p = 0.029 | 0.092** (0.028)<br>p = 0.003 | 0.137*** (0.032)<br>p = 0.0002 |
| Amyloid group<br>(ref = amyloid-) | 0.040 (0.026)<br>p = 0.127 | 0.095** (0.028)<br>p = 0.002 | 0.072* (0.028)<br>p = 0.012 | 0.078* (0.032)<br>p = 0.019 |
| Age<br>× amyloid group | 0.009** (0.003)<br>p = 0.003 | 0.005 (0.003)<br>p = 0.118 | 0.010** (0.003)<br>p = 0.003 | 0.009* (0.004)<br>p = 0.014 |

*Note:*
\* $p < 0.05$ ; \*\* $p < 0.01$ ; \*\*\* $p < 0.001$

## Appendix F. Peak tables

MNI coordinates of peak voxels and local maxima within significant clusters were obtained using `atlasreader` [64]. We transformed these MNI coordinates into Talairach coordinates [65] using an inhouse Python implementation of the coordinate look-up procedure implemented in BioImage Suite Web, which is based on a non-linear mapping between a digitized Talairach atlas and the MNI template [66]. We performed a 9 mm-wide cube range search using the Talairach Client (http://www.talairach.org/client.html) to obtain anatomical labels for each peak and subpeak. The label with the most hits in the cube was chosen as the corresponding anatomical label. For top hits that were not assigned a Brodmann area (BA) in the Talairach client output, if there was another hit with the same anatomical label as the top hit, we report their BA where applicable. All label and BA assignments were confirmed via visual inspection and corrected as necessary. (Sub)Peaks are sorted by the laterality (Bilateral, Left hemisphere, or Right hemisphere) of the cluster they belong to, then the brain region (Frontal, Parietal, Temporal, Occipital, Cingulate, Midbrain, Pons, Subcortical, Cerebellum, or a combination thereof) the cluster falls into. We omitted subpeaks with repeated labels within a cluster, and display only the one with the largest T-value. *Abbreviations*: BA = Brodmann area, L = Left, R = Right.

**Table F.1:** Cluster peaks and subpeaks for the age × amyloid status interaction.

| Label | Volume<br>(mm<sup>3</sup>) | MNI ($x, y, z$) | T-value |
| --- | --- | --- | --- |
| **Bilateral** |  |  |  |
| *Cingulate* |  |  |  |
| L Cingulate Gyrus BA 31 | 783 | (-9, -52, 25) | 4.56 |
| R Cingulate Gyrus BA 31 |  | (2, -47, 29) | 3.83 |
| **Left hemisphere** |  |  |  |
| *Frontal* |  |  |  |
| Middle Frontal Gyrus BA 10 | 604 | (-34, 50, 8) | 4.20 |
| Middle Frontal Gyrus BA 6 | 993 | (-42, 4, 46) | 5.09 |
| Precentral Gyrus BA 6 |  | (-40, -1, 32) | 4.38 |
| *Frontal, Cingulate* |  |  |  |
| Superior Frontal Gyrus BA 9 | 928 | (-15, 37, 42) | 4.29 |
| Anterior Cingulate BA 32 |  | (-15, 35, 17) | 4.24 |
| *Frontal, Parietal* |  |  |  |
| Insula BA 13 | 1588 | (-39, -10, 18) | 3.88 |
| Postcentral Gyrus BA 43 |  | (-52, -15, 19) | 4.56 |
| *Frontal, Temporal* |  |  |  |
| Frontal Sub-Gyrus | 845 | (-47, 10, 11) | 4.63 |
| Superior Temporal Gyrus BA 22 |  | (-51, 5, -1) | 4.78 |
| *Occipital* |  |  |  |
| Cuneus BA 18 | 670 | (0, -87, 18) | 5.13 |
|  | 640 | (-5, -97, 1) | 4.89 |
| **Right hemisphere** |  |  |  |
| *Cingulate* |  |  |  |
| Anterior Cingulate BA 32 | 409 | (7, 43, 6) | 5.06 |
| *Frontal* |  |  |  |
| Frontal Sub-Gyrus | 1320 | (24, 41, -6) | 4.87 |
| Inferior Frontal Gyrus BA 47 |  | (34, 32, -9) | 5.75 |
| Superior Frontal Gyrus BA 6/8 | 614 | (12, 25, 56) | 4.66 |
| *Occipital* |  |  |  |
| Cuneus BA 17 | 560 | (15, -92, 7) | 4.58 |
| Middle Occipital Gyrus BA 18 |  | (20, -94, 18) | 4.00 |
| *Parietal* |  |  |  |
| Inferior Parietal Lobule BA 40 | 793 | (57, -36, 25) | 4.41 |
| Postcentral Gyrus BA 2 |  | (54, -25, 32) | 3.97 |
| *Temporal* |  |  |  |
| Middle Temporal Gyrus BA 37 | 485 | (56, -47, -16) | 4.58 |
| *Temporal, Occipital* |  |  |  |
| Middle Temporal Gyrus BA 39 | 1326 | (44, -72, 14) | 4.23 |
| Middle Occipital Gyrus BA 19 |  | (45, -77, -8) | 5.21 |

**Table F.2:** Cluster peaks and subpeaks for the main effect of age among amyloid+ individuals.

| Label | Volume<br>(mm<sup>3</sup>) | MNI ($x, y, z$) | T-value |
| --- | --- | --- | --- |
| **Left hemisphere** |  |  |  |
| *Subcortical* |  |  |  |
| Putamen | 515 | (-20, 8, 3) | 4.30 |
| **Right hemisphere** |  |  |  |
| *Frontal* |  |  |  |
| Inferior Frontal Gyrus BA 47 | 415 | (35, 31, -9) | 5.47 |
| *Occipital* |  |  |  |
| Middle Occipital Gyrus BA 19 | 557 | (45, -77, -8) | 5.75 |
| *Subcortical* |  |  |  |
| Putamen | 2660 | (31, 4, 1) | 5.38 |

**Table F.3:**
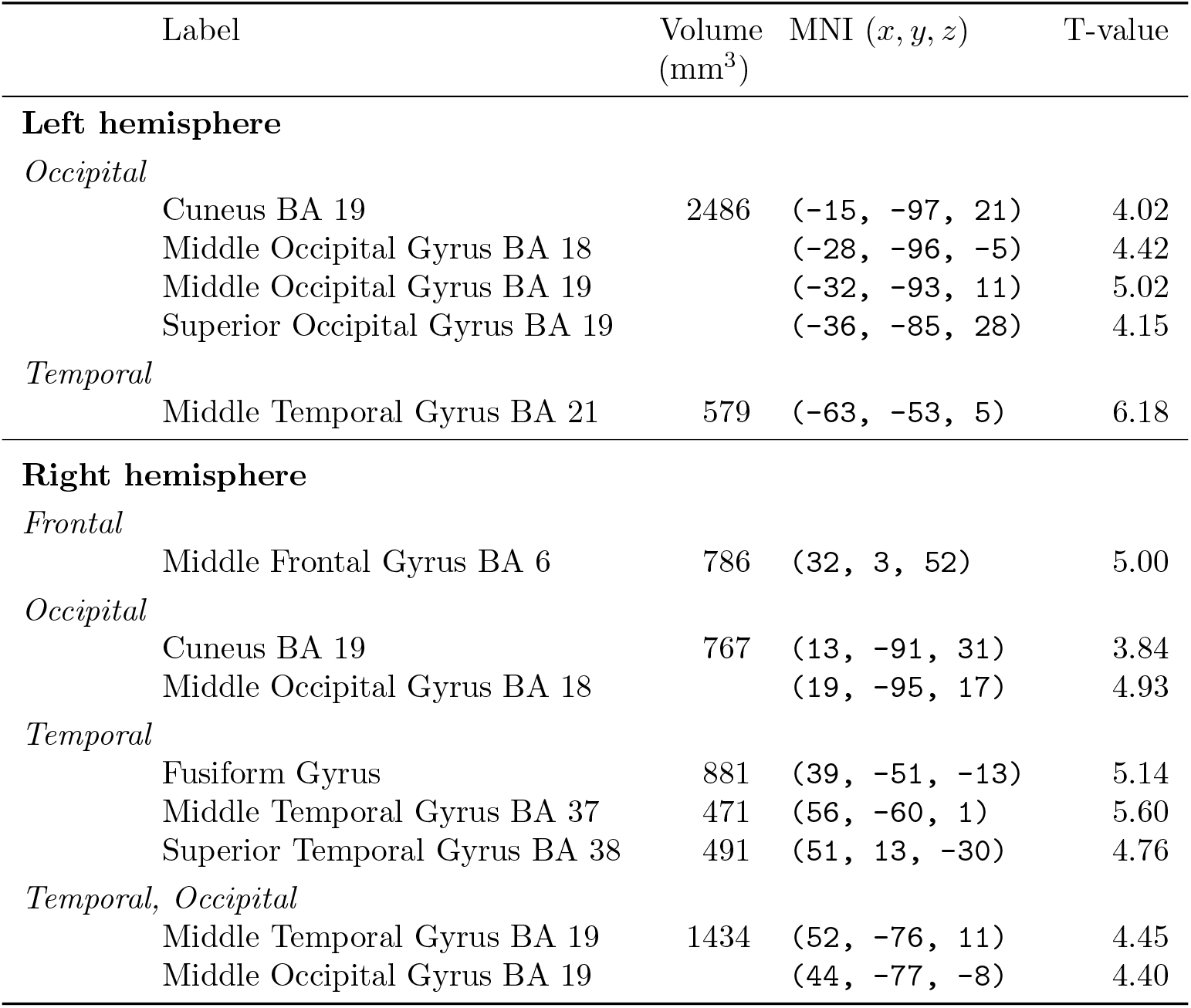
Cluster peaks and subpeaks for the main effect of amyloid status. Positive T-values indicate that ^18^F-AV-1451 SUVR is greater among amyloid+ compared to amyloid–individuals.

**Table F.4:**
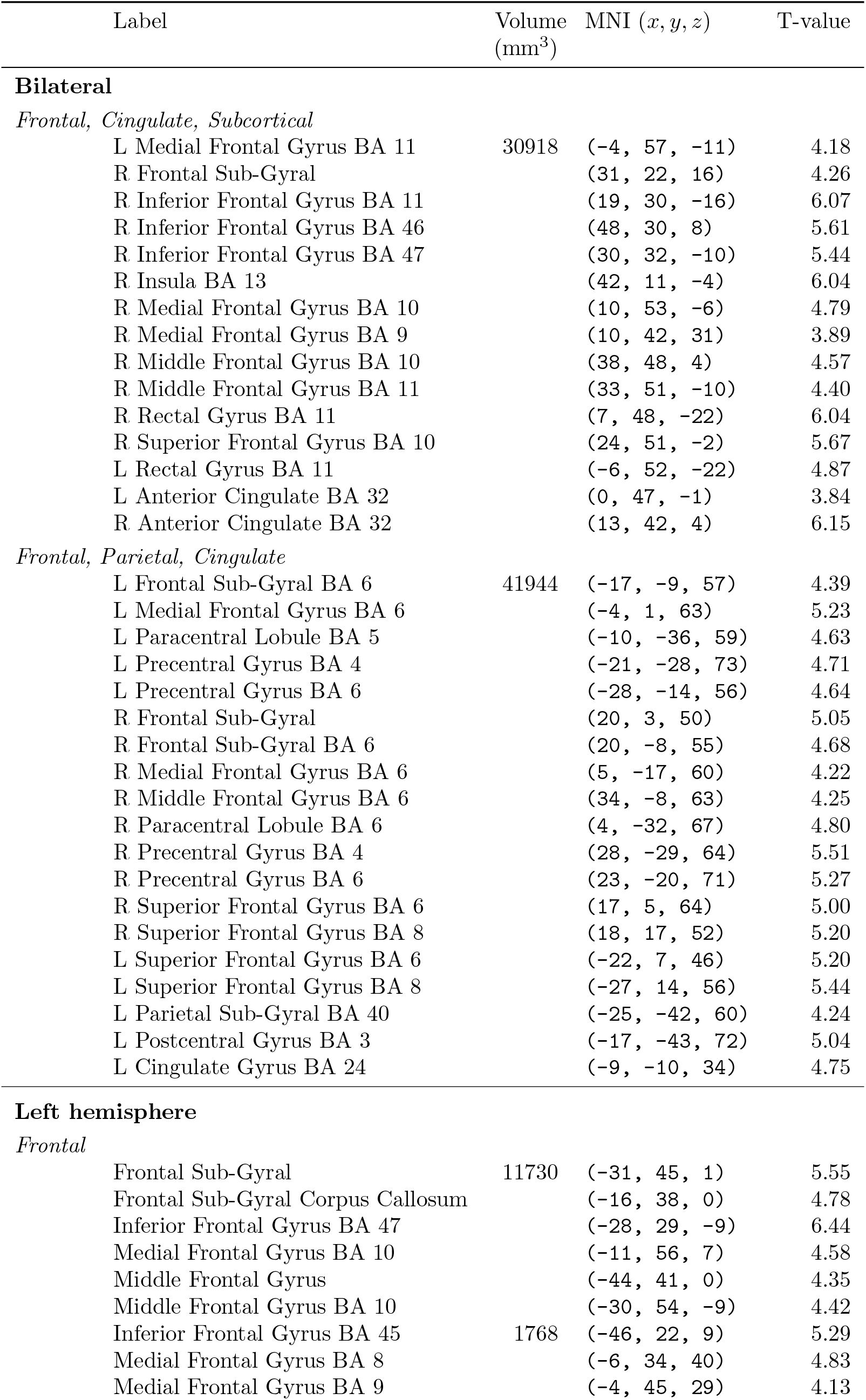
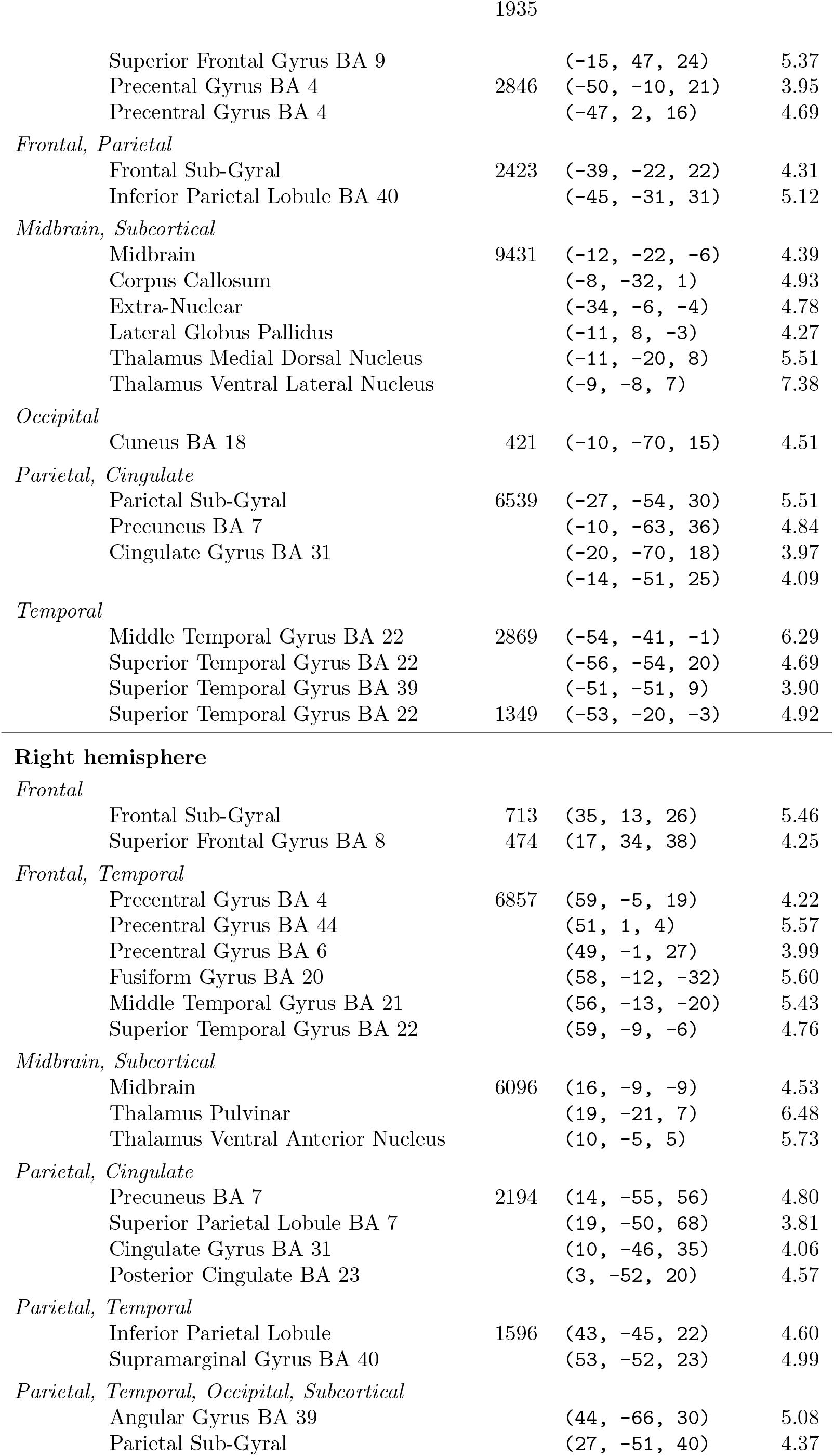
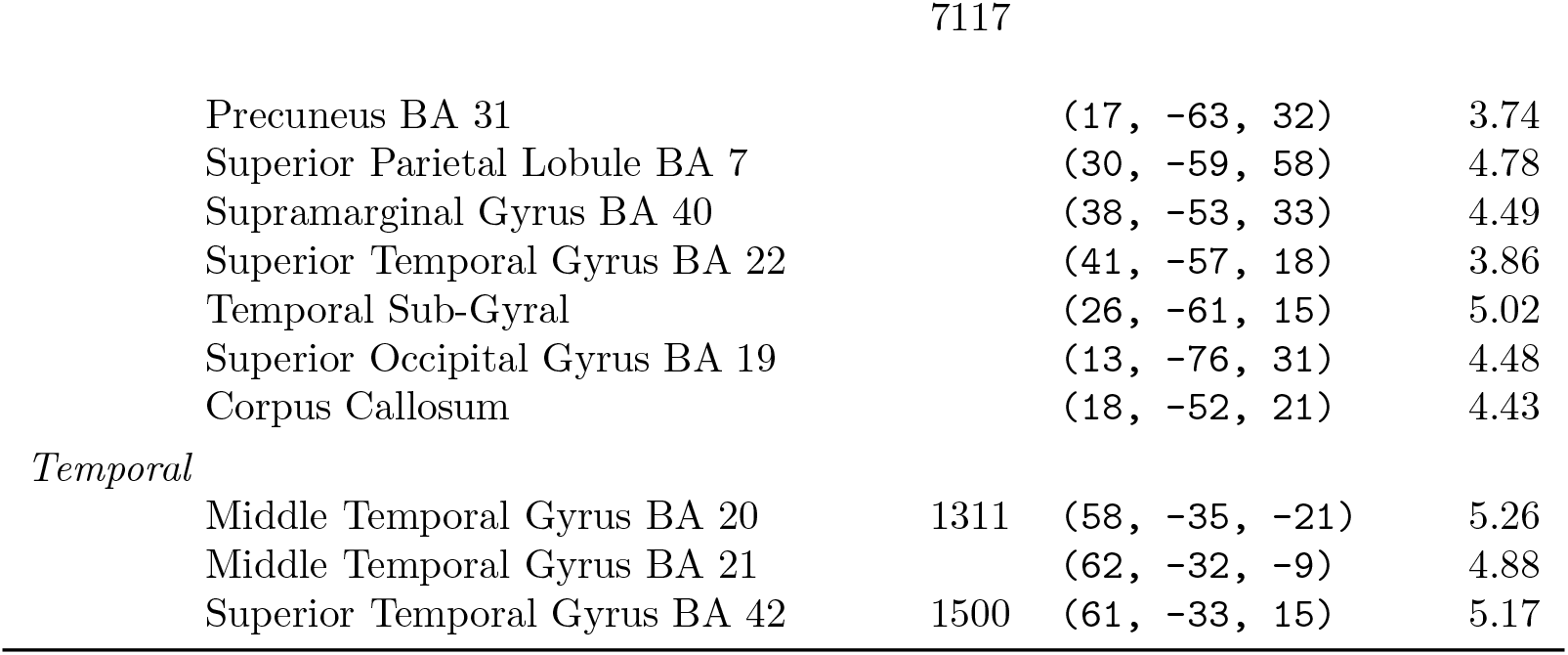
Cluster peaks and subpeaks for the main effect of sex. Positive T-values indicate that ^18^F-AV-1451 SUVR is greater among men compared to women.

**Table F.5:** Cluster peaks and subpeaks for the main effect of race. Positive T-values indicate that ^18^F-AV-1451 SUVR is greater among black compared to non-black individuals.

| Label | Volume<br>(mm<sup>3</sup>) | MNI ($x, y, z$) | T-value |
| --- | --- | --- | --- |
| **Bilateral** |  |  |  |
| *Frontal* |  |  |  |
| R Middle Frontal Gyrus BA 6 | 13235 | (33, 2, 58) | 5.77 |
| R Superior Frontal Gyrus BA 6 |  | (8, 22, 61) | 5.31 |
| R Superior Frontal Gyrus BA 8 |  | (20, 19, 50) | 4.90 |
| L Superior Frontal Gyrus BA 8 |  | (-10, 37, 54) | 4.66 |
| L Superior Frontal Gyrus BA 9 |  | (-12, 60, 33) | 3.74 |
| **Left hemisphere** |  |  |  |
| *Cerebellum* |  |  |  |
| Anterior Lobe | 826 | (-17, -42, -37) | 4.23 |
|  | 420 | (-27, -47, -39) | 4.77 |
| *Frontal* |  |  |  |
| Middle Frontal Gyrus BA 46 | 1371 | (-41, 28, 22) | 4.29 |
| Middle Frontal Gyrus BA 9 |  | (-41, 35, 32) | 4.94 |
| Middle Frontal Gyrus BA 6 | 959 | (-34, 9, 62) | 4.88 |
| *Occipital* |  |  |  |
| Cuneus BA 17 | 1358 | (-19, -73, 22) | 5.14 |
| Cuneus BA 18 |  | (-3, -74, 14) | 3.78 |
|  | 755 | (-8, -74, 2) | 4.13 |
| Lingual Gyrus BA 18 |  | (-17, -74, -9) | 4.76 |
| Middle Occipital Gyrus BA 18 | 1855 | (-32, -86, -1) | 4.38 |
| Superior Occipital Gyrus BA 19 |  | (-31, -84, 26) | 4.54 |
| *Parietal* |  |  |  |
| Precuneus BA 7 | 2368 | (-5, -77, 42) | 5.09 |
| *Subcortical* |  |  |  |
| Thalamus Medial Dorsal Nucleus | 862 | (-11, -21, 10) | 4.19 |
| Thalamus Pulvinar |  | (-23, -30, 6) | 3.99 |
| **Right hemisphere** |  |  |  |
| *Frontal* |  |  |  |
| Precentral Gyrus BA 6 | 1236 | (27, -20, 71) | 4.95 |
| *Occipital* |  |  |  |
| Cuneus BA 17 | 2669 | (16, -89, 2) | 4.66 |
| Lingual Gyrus BA 17 |  | (8, -91, -10) | 4.92 |
| Lingual Gyrus BA 18 |  | (9, -79, -2) | 3.93 |
| Lingual Gyrus BA 19 |  | (18, -67, -4) | 5.11 |
| Middle Occipital Gyrus BA 19 | 506 | (28, -86, 12) | 4.04 |
| *Temporal* |  |  |  |
| Inferior Temporal Gyrus BA 20 | 890 | (60, -17, -30) | 4.36 |
| Middle Temporal Gyrus BA 21 |  | (59, -13, -15) | 5.49 |
| Middle Temporal Gyrus BA 20 | 514 | (61, -39, -15) | 4.94 |
| Parahippocampal Gyrus BA 27 | 531 | (22, -30, -10) | 4.70 |
| *Temporal, Midbrain* |  |  |  |
| Parahippocampal Gyrus BA 28 | 415 | (23, -20, -14) | 3.73 |
| Midbrain |  | (9, -7, -9) | 4.28 |

## Appendix G. Cognition and tau accumulation

**Table G.1:** Linear mixed effects models of the relationship between entorhinal ^18^F-AV-1451 SUVR and cognition. Estimated fixed effects are reported along with their standard errors in parentheses.

|  | *Dependent variable: Cognitive testing performance (z-score)* |  |  |  |  |  |
| --- | --- | --- | --- | --- | --- | --- |
|  | Memory | Attention | Executive Function | Fluency | Visuospatial Processing | Processing Speed |
| Intercept | 0.126 (0.125)<br>p = 0.312 | 0.119 (0.095)<br>p = 0.213 | 0.014 (0.099)<br>p = 0.891 | 0.062 (0.125)<br>p = 0.619 | -0.049 (0.092)<br>p = 0.597 | 0.019 (0.117)<br>p = 0.869 |
| Age at PET scan | -0.029* (0.012)<br>p = 0.021 | -0.023* (0.011)<br>p = 0.037 | -0.021 (0.011)<br>p = 0.058 | -0.028* (0.013)<br>p = 0.038 | -0.022* (0.010)<br>p = 0.030 | -0.055*** (0.013)<br>p = 0.0001 |
| Sex<br>(ref = female) | -0.082 (0.217)<br>p = 0.708 | -0.185 (0.188)<br>p = 0.331 | -0.064 (0.195)<br>p = 0.745 | 0.022 (0.239)<br>p = 0.928 | 0.151 (0.163)<br>p = 0.361 | -0.450 (0.230)<br>p = 0.057 |
| Education (years) | -0.066 (0.051)<br>p = 0.205 | 0.016 (0.045)<br>p = 0.717 | -0.006 (0.046)<br>p = 0.896 | -0.052 (0.056)<br>p = 0.354 | -0.005 (0.038)<br>p = 0.889 | 0.148** (0.055)<br>p = 0.010 |
| Amyloid group<br>(ref = amyloid-) | 0.131 (0.308)<br>p = 0.673 | 0.118 (0.238)<br>p = 0.623 | -0.294 (0.247)<br>p = 0.241 | -0.137 (0.310)<br>p = 0.661 | 0.030 (0.226)<br>p = 0.896 | 0.021 (0.291)<br>p = 0.943 |
| Entorhinal SUVR | -0.769 (0.942)<br>p = 0.419 | -0.951 (0.714)<br>p = 0.190 | 0.321 (0.746)<br>p = 0.670 | -0.503 (0.946)<br>p = 0.598 | 0.399 (0.688)<br>p = 0.565 | -0.242 (0.885)<br>p = 0.786 |
| Time from PET | -0.023*** (0.006)<br>p = 0.0001 | -0.020*** (0.006)<br>p = 0.001 | -0.016*** (0.004)<br>p = 0.0004 | -0.011* (0.005)<br>p = 0.016 | -0.024*** (0.005)<br>p = 0.000002 | -0.097*** (0.010)<br>p = 0.000 |
| Amyloid group<br>× time | 0.007 (0.014)<br>p = 0.646 | -0.016 (0.015)<br>p = 0.271 | -0.019 (0.010)<br>p = 0.056 | 0.023* (0.011)<br>p = 0.047 | 0.006 (0.011)<br>p = 0.588 | -0.038 (0.026)<br>p = 0.147 |
| Entorhinal SUVR<br>× time | -0.086* (0.039)<br>p = 0.029 | 0.017 (0.033)<br>p = 0.611 | 0.035 (0.023)<br>p = 0.128 | -0.037 (0.026)<br>p = 0.161 | 0.045 (0.027)<br>p = 0.095 | -0.001 (0.073)<br>p = 0.988 |

Note:
\*p<0.05; \*\*p<0.01; \*\*\*p<0.001

**Table G.2:** Linear mixed effects models of the relationship between hippocampus ^18^F-AV-1451 SUVR and cognition. Estimated fixed effects are reported along with their standard errors in parentheses.

|  | *Dependent variable: Cognitive testing performance (z-score)* |  |  |  |  |  |
| --- | --- | --- | --- | --- | --- | --- |
|  | Memory | Attention | Executive Function | Fluency | Visuospatial Processing | Processing Speed |
| Intercept | 0.126 (0.121)<br>p = 0.300 | 0.122 (0.095)<br>p = 0.200 | 0.022 (0.099)<br>p = 0.828 | 0.059 (0.121)<br>p = 0.627 | -0.044 (0.092)<br>p = 0.628 | 0.020 (0.116)<br>p = 0.862 |
| Age at PET scan | -0.028* (0.012)<br>p = 0.023 | -0.024* (0.010)<br>p = 0.028 | -0.022 (0.011)<br>p = 0.053 | -0.026* (0.013)<br>p = 0.046 | -0.022* (0.010)<br>p = 0.025 | -0.055*** (0.013)<br>p = 0.0001 |
| Sex<br>(ref = female) | -0.113 (0.213)<br>p = 0.597 | -0.186 (0.187)<br>p = 0.324 | -0.053 (0.194)<br>p = 0.788 | 0.008 (0.226)<br>p = 0.972 | 0.186 (0.161)<br>p = 0.255 | -0.453 (0.229)<br>p = 0.055 |
| Education (years) | -0.051 (0.052)<br>p = 0.333 | 0.034 (0.046)<br>p = 0.468 | 0.001 (0.048)<br>p = 0.980 | -0.048 (0.056)<br>p = 0.392 | -0.006 (0.040)<br>p = 0.890 | 0.154** (0.056)<br>p = 0.010 |
| Amyloid group<br>(ref = amyloid-) | 0.071 (0.293)<br>p = 0.810 | 0.056 (0.231)<br>p = 0.810 | -0.281 (0.242)<br>p = 0.252 | -0.202 (0.293)<br>p = 0.494 | 0.052 (0.221)<br>p = 0.817 | 0.005 (0.284)<br>p = 0.986 |
| Hippocampus SUVR | -0.833* (0.405)<br>p = 0.046 | -0.456 (0.316)<br>p = 0.156 | -0.207 (0.332)<br>p = 0.537 | -0.766 (0.405)<br>p = 0.065 | 0.153 (0.304)<br>p = 0.618 | -0.174 (0.392)<br>p = 0.661 |
| Time from PET | -0.024*** (0.006)<br>p = 0.0001 | -0.020*** (0.006)<br>p = 0.001 | -0.014** (0.005)<br>p = 0.002 | -0.013*** (0.004)<br>p = 0.001 | -0.023*** (0.005)<br>p = 0.00001 | -0.097*** (0.010)<br>p = 0.000 |
| Amyloid group<br>× time | 0.002 (0.015)<br>p = 0.917 | -0.015 (0.015)<br>p = 0.310 | -0.018 (0.011)<br>p = 0.109 | 0.017* (0.009)<br>p = 0.050 | 0.008 (0.011)<br>p = 0.467 | -0.038 (0.026)<br>p = 0.143 |
| Hippocampus SUVR<br>× time | -0.041* (0.017)<br>p = 0.018 | 0.010 (0.014)<br>p = 0.459 | 0.003 (0.010)<br>p = 0.741 | -0.028*** (0.008)<br>p = 0.001 | 0.014 (0.011)<br>p = 0.212 | 0.0005 (0.030)<br>p = 0.988 |

Note:
\*p<0.05; \*\*p<0.01; \*\*\*p<0.001

**Table G.3:** Linear mixed effects models of the relationship between inferior temporal gyrus ^18^F-AV-1451 SUVR and cognition. Estimated fixed effects are reported along with their standard errors in parentheses.

|  | *Dependent variable: Cognitive testing performance (z-score)* |  |  |  |  |  |
| --- | --- | --- | --- | --- | --- | --- |
|  | Memory | Attention | Executive Function | Fluency | Visuospatial Processing | Processing Speed |
| Intercept | 0.122 (0.124)<br>p = 0.327 | 0.118 (0.092)<br>p = 0.202 | 0.015 (0.098)<br>p = 0.881 | 0.061 (0.124)<br>p = 0.624 | -0.048 (0.092)<br>p = 0.599 | 0.018 (0.116)<br>p = 0.878 |
| Age at PET scan | -0.029* (0.012)<br>p = 0.021 | -0.024* (0.010)<br>p = 0.022 | -0.021 (0.011)<br>p = 0.056 | -0.029* (0.013)<br>p = 0.033 | -0.021* (0.010)<br>p = 0.030 | -0.055*** (0.013)<br>p = 0.0001 |
| Sex<br>(ref = female) | -0.096 (0.225)<br>p = 0.670 | -0.042 (0.189)<br>p = 0.825 | 0.045 (0.195)<br>p = 0.817 | 0.038 (0.248)<br>p = 0.880 | 0.200 (0.162)<br>p = 0.224 | -0.405 (0.237)<br>p = 0.095 |
| Education (years) | -0.062 (0.051)<br>p = 0.237 | 0.023 (0.043)<br>p = 0.598 | -0.003 (0.045)<br>p = 0.950 | -0.052 (0.057)<br>p = 0.364 | 0.0001 (0.038)<br>p = 0.998 | 0.149** (0.054)<br>p = 0.009 |
| Amyloid group<br>(ref = amyloid-) | 0.284 (0.338)<br>p = 0.406 | 0.316 (0.251)<br>p = 0.215 | -0.097 (0.267)<br>p = 0.718 | -0.052 (0.340)<br>p = 0.881 | 0.095 (0.250)<br>p = 0.706 | 0.085 (0.319)<br>p = 0.790 |
| ITG SUVR | -1.647 (1.410)<br>p = 0.249 | -2.451* (1.046)<br>p = 0.024 | -1.642 (1.118)<br>p = 0.149 | -1.021 (1.422)<br>p = 0.477 | -0.346 (1.044)<br>p = 0.742 | -0.734 (1.334)<br>p = 0.585 |
| Time from PET | -0.024*** (0.006)<br>p = 0.00005 | -0.020*** (0.006)<br>p = 0.001 | -0.015*** (0.004)<br>p = 0.001 | -0.011* (0.005)<br>p = 0.023 | -0.023*** (0.005)<br>p = 0.00001 | -0.097*** (0.010)<br>p = 0.000 |
| Amyloid group<br>× time | 0.017 (0.016)<br>p = 0.300 | -0.015 (0.016)<br>p = 0.344 | -0.018 (0.011)<br>p = 0.112 | 0.028* (0.013)<br>p = 0.037 | 0.003 (0.013)<br>p = 0.816 | -0.040 (0.028)<br>p = 0.157 |
| ITG SUVR<br>× time | -0.132* (0.066)<br>p = 0.048 | -0.010 (0.062)<br>p = 0.879 | 0.023 (0.045)<br>p = 0.610 | -0.060 (0.051)<br>p = 0.239 | 0.056 (0.051)<br>p = 0.277 | 0.029 (0.122)<br>p = 0.815 |

Note:
\*p<0.05; \*\*p<0.01; \*\*\*p<0.001

**Table G.4:** Linear mixed effects models of the relationship between entorhinal ^18^F-AV-1451 SUVR and California Verbal Learning Test (CVLT) *z*-scores. Estimated fixed effects are reported along with their standard errors in parentheses.

|  | *Dependent variable: CVLT ($z$-score)* |  |
| --- | --- | --- |
|  | Immediate recall | Long-delay free recall |
| Intercept | 0.105 (0.122)<br>p = 0.391 | 0.146 (0.132)<br>p = 0.269 |
| Age at PET scan | -0.032* (0.012)<br>p = 0.011 | -0.027* (0.013)<br>p = 0.042 |
| Sex<br>(ref = female) | -0.101 (0.214)<br>p = 0.641 | -0.037 (0.230)<br>p = 0.875 |
| Education (years) | -0.089 (0.051)<br>p = 0.084 | -0.047 (0.054)<br>p = 0.397 |
| Amyloid group<br>(ref = amyloid-) | 0.127 (0.302)<br>p = 0.677 | 0.122 (0.326)<br>p = 0.711 |
| Entorhinal SUVR | -0.645 (0.924)<br>p = 0.489 | -0.893 (0.996)<br>p = 0.375 |
| Time from PET | -0.024*** (0.006)<br>p = 0.0001 | -0.022*** (0.006)<br>p = 0.0005 |
| Amyloid group<br>× time | 0.011 (0.015)<br>p = 0.480 | 0.001 (0.015)<br>p = 0.965 |
| Entorhinal SUVR<br>× time | -0.078 (0.041)<br>p = 0.060 | -0.095* (0.042)<br>p = 0.024 |
| *Note:* *p<0.05; **p<0.01; ***p<0.001 |  |  |

## References

1. Goedert M, Spillantini MG, Jakes R, Rutherford D, Crowther RA. Multiple isoforms of human microtubule-associated protein tau: sequences and localization in neurofibrillary tangles of Alzheimer’s disease. Neuron 1989;3(4):519–26. doi:10.1016/0896-6273(89)90210-9.

2. Nelson PT, Alafuzoff I, Bigio EH, Bouras C, Braak H, Cairns NJ, Castellani RJ, Crain BJ, Davies P, Del Tredici K, Duyckaerts C, Frosch MP, Haroutunian V, Hof PR, Hulette CM, Hyman BT, Iwatsubo T, Jellinger KA, Jicha GA, Kövari E, Kukull WA, Leverenz JB, Love S, Mackenzie IR, Mann DM, Masliah E, McKee AC, Montine TJ, Morris JC, Schneider JA, Sonnen JA, Thal DR, Trojanowski JQ, Troncoso JC, Wisniewski T, Woltjer RL, Beach TG. Correlation of Alzheimer disease neuropathologic changes with cognitive status: a review of the literature. Journal of Neuropathology and Experimental Neurology 2012;71(5):362–81. doi:10.1097/NEN.0b013e31825018f7.

3. Crary JF, Trojanowski JQ, Schneider JA, Abisambra JF, Abner EL, Alafuzoff I, Arnold SE, Attems J, Beach TG, Bigio EH, Cairns NJ, Dickson DW, Gearing M, Grinberg LT, Hof PR, Hyman BT, Jellinger K, Jicha GA, Kovacs GG, Knopman DS, Kofler J, Kukull WA, Mackenzie IR, Masliah E, McKee A, Montine TJ, Murray ME, Neltner JH, Santa-Maria I, Seeley WW, Serrano-Pozo A, Shelanski ML, Stein T, Takao M, Thal DR, Toledo JB, Troncoso JC, Vonsattel JP, White CL, Wisniewski T, Woltjer RL, Yamada M, Nelson PT. Primary age-related tauopathy (PART): a common pathology associated with human aging. Acta Neuropathologica, 2014;128(6):755–66. doi:10.1007/s00401-014-1349-0.

4. Duyckaerts C, Braak H, Brion JP, Bueé L, Del Tredici K, Goedert M, Halliday G, Neumann M, Spillantini MG, Tolnay M, Uchihara T. PART is part of Alzheimer disease. Acta Neuropathologica 2015;129(5):749–56. doi:10.1007/s00401-015-1390-7.

5. Jack CR, Knopman DS, Jagust WJ, Petersen RC, Weiner MW, Aisen PS, Shaw LM, Vemuri P, Wiste HJ, Weigand SD, Lesnick TG, Pankratz VS, Donohue MC, Trojanowski JQ. Tracking pathophysiological processes in Alzheimer’s disease: an updated hypothetical model of dynamic biomarkers. The Lancet Neurology 2013;12(2):207–16. doi:10.1016/S1474-4422(12)70291-0.

6. Braak H, Braak E. Neuropathological stageing of Alzheimer-related changes. Acta Neuropathologica 1991;82(4):239–59. doi:10.1007/BF00308809.

7. Brier MR, Gordon B, Friedrichsen K, McCarthy J, Stern A, Christensen J, Owen C, Aldea P, Su Y, Hassenstab J, Cairns NJ, Holtzman DM, Fagan AM, Morris JC, Benzinger TLS, Ances BM. Tau and A*β* imaging, CSF measures, and cognition in Alzheimer’s disease. Science Translational Medicine 2016;8(338):338ra66. doi:10.1126/scitranslmed.aaf2362.

8. Maass A, Landau S, Baker SL, Horng A, Lockhart SN, La Joie R, Rabinovici GD, Jagust WJ, Alzheimer’s Disease Neuroimaging Initiative. Comparison of multiple tau-PET measures as biomarkers in aging and Alzheimer’s disease. NeuroImage 2017;157:448–63. doi:10.1016/j.neuroimage.2017.05.058.

9. Pontecorvo MJ, Devous MD, Navitsky M, Lu M, Salloway S, Schaerf FW, Jennings D, Arora AK, McGeehan A, Lim NC, Xiong H, Joshi AD, Siderowf A, Mintun MA, investigators FAA. Relationships between flortaucipir PET tau binding and amyloid burden, clinical diagnosis, age and cognition. Brain 2017;140(3):748–63. doi:10.1093/brain/aww334.

10. Wang L, Benzinger TL, Su Y, Christensen J, Friedrichsen K, Aldea P, McConathy J, Cairns NJ, Fagan AM, Morris JC, Ances BM. Evaluation of tau imaging in staging Alzheimer disease and revealing interactions between *β*-amyloid and tauopathy. JAMA Neurology 2016;73(9):1070–7. doi:10.1001/jamaneurol.2016.2078.

11. Das SR, Xie L, Wisse LEM, Ittyerah R, Tustison NJ, Dickerson BC, Yushkevich PA, Wolk DA, Alzheimer’s Disease Neuroimaging Initiative. Longitudinal and cross-sectional structural magnetic resonance imaging correlates of AV-1451 uptake. Neurobiology of Aging 2018;66:49–58. doi:10.1016/j.neurobiolaging.2018.01.024.

12. Iaccarino L, Tammewar G, Ayakta N, Baker SL, Bejanin A, Boxer AL, Gorno-Tempini ML, Janabi M, Kramer JH, Lazaris A, Lockhart SN, Miller BL, Miller ZA, O’Neil JP, Ossenkoppele R, Rosen HJ, Schonhaut DR, Jagust WJ, Rabinovici GD. Local and distant relationships between amyloid, tau and neurodegeneration in Alzheimer’s disease. NeuroImage: Clinical 2018;17:452–64. doi:10.1016/j.nicl.2017.09.016.

13. Aschenbrenner AJ, Gordon BA, Benzinger TLS, Morris JC, Hassenstab JJ. Influence of tau PET, amyloid PET, and hippocampal volume on cognition in Alzheimer disease. Neurology 2018;91(9):e859–66. doi:10.1212/WNL.0000000000006075.

14. Koychev I, Gunn RN, Firouzian A, Lawson J, Zamboni G, Ridha B, Sahakian BJ, Rowe JB, Thomas A, Rochester L, Ffytche D, Howard R, Zetterberg H, MacKay C, Lovestone S, Deep and Frequent Phenotyping study team. PET tau and amyloid-*β* burden in mild Alzheimer’s disease: divergent relationship with age, cognition, and cerebrospinal fluid biomarkers. Journal of Alzheimer’s Disease 2017;60(1):283–93. doi:10.3233/JAD-170129.

15. Pontecorvo MJ, Devous MD, Kennedy I, Navitsky M, Lu M, Galante N, Salloway S, Doraiswamy PM, Southekal S, Arora AK, McGeehan A, Lim NC, Xiong H, Truocchio SP, Joshi AD, Shcherbinin S, Teske B, Fleisher AS, Mintun MA. A multicentre longitudinal study of flortaucipir (^18^F) in normal ageing, mild cognitive impairment and Alzheimer’s disease dementia. Brain 2019;:1–13doi:10.1093/brain/awz090.

16. Sepulcre J, Schultz AP, Sabuncu M, Gomez-Isla T, Chhatwal J, Becker A, Sperling R, Johnson KA. In vivo tau, amyloid, and gray matter profiles in the aging brain. Journal of Neuroscience 2016;36(28):7364–74. doi:10.1523/JNEUROSCI.0639-16.2016.

17. LaPoint MR, Chhatwal JP, Sepulcre J, Johnson KA, Sperling RA, Schultz AP. The association between tau PET and retrospective cortical thinning in clinically normal elderly. NeuroImage 2017;157(May):612–22. doi:10.1016/j.neuroimage.2017.05.049.

18. Maass A, Lockhart SN, Harrison TM, Bell RK, Mellinger T, Swinnerton K, Baker SL, Rabinovici GD, Jagust WJ. Entorhinal tau pathology, episodic memory decline, and neurodegeneration in aging. The Journal of Neuroscience 2018;38(3):530–43. doi:10.1523/JNEUROSCI.2028-17.2017.

19. Scholl M, Lockhart SN, Schonhaut DR, O’Neil JP, Janabi M, Ossenkoppele R, Baker SL, Vogel JW, Faria J, Schwimmer HD, Rabinovici GD, Jagust WJ. PET imaging of tau deposition in the aging human brain. Neuron 2016;89(5):971–82. doi:10.1016/j.neuron.2016.01.028.

20. Schultz SA, Gordon BA, Mishra S, Su Y, Perrin RJ, Cairns NJ, Morris JC, Ances BM, Benzinger TLS. Widespread distribution of tauopathy in preclinical Alzheimer’s disease. Neurobiology of Aging 2018;72:177–85. doi:10.1016/j.neurobiolaging.2018.08.022.

21. Sperling RA, Mormino EC, Schultz AP, Betensky RA, Papp KV, Amariglio RE, Hanseeuw BJ, Buckley R, Chhatwal J, Hedden T, Marshall GA, Quiroz YT, Donovan NJ, Jackson J, Gatchel JR, Rabin JS, Jacobs H, Yang H, Properzi M, Kirn DR, Rentz DM, Johnson KA. The impact of amyloid-beta and tau on prospective cognitive decline in older individuals. Annals of Neurology 2019;85(2):181–93. doi:10.1002/ana.25395.

22. Morris JC. The Clinical Dementia Rating (CDR): Current version and scoring rules. 1993.

23. Fuld PA. Psychological testing in the differential diagnosis of the dementias. In: Katzman R, Terry RD, Bick KL, eds. Alzheimer’s disease: Senile dementia and related disorders. New York, NY: Raven Press; 1978:185–93.

24. Doshi J, Erus G, Ou Y, Resnick SM, Gur RC, Gur RE, Satterthwaite TD, Furth S, Davatzikos C. MUSE: MUlti-atlas region Segmentation utilizing Ensembles of registration algorithms and parameters, and locally optimal atlas selection. NeuroImage 2016;127(2016):186–95. doi:10.1016/j.neuroimage.2015.11.073.

25. Jack CR, Twomey K, Zinsmeister AR, Sharbrough FW, Petersen C, Cascino GD. Anterior temporal lobes and hippocampal formations: normative volumetric measurements from MR images in young adults. Radiology 1989;172:549–54.

26. Thomas BA, Erlandsson K, Modat M, Thurfjell L, Vandenberghe R, Ourselin S, Hutton BF. The importance of appropriate partial volume correction for PET quantification in Alzheimer’s disease. European Journal of Nuclear Medicine and Molecular Imaging 2011;38(6):1104–19. doi:10.1007/s00259-011-1745-9.

27. Baker SL, Maass A, Jagust WJ. Considerations and code for partial volume correcting [^18^F]-AV-1451 tau PET data. Data in Brief 2017;15:648–57. doi:10.1016/j.dib.2017.10.024.

28. Diedrichsen J, Balsters JH, Flavell J, Cussans E, Ramnani N. A probabilistic MR atlas of the human cerebellum. NeuroImage 2009;46(1):39–46. doi:10.1016/j.neuroimage.2009.01.045.

29. Delis DC, Kramer JH, Kaplan E, Ober BA. The California Verbal Learning Test. San Antonio, TX: Psychological Corporation; 1987.

30. Reitan RM. Trail making test: Manual for administration and scoring. Tucson, AZ: Reitan Neuropsychological Laboratory; 1992.

31. Wechsler D. Wechsler adult intelligence scale-revised. Revised ed.; San Antonio, TX: Psychological Corporation; 1981.

32. Newcombe F. Missile wounds of the brain: a study of psychological deficits. Oxford: Oxford University Press; 1969.

33. Benton AL. Differential behavioral effects in frontal lobe disease. Neuropsychologia 1968;6(1):53–60. doi:10.1016/0028-3932(68)90038-9.

34. Wilson JR, De Fries JC, Mc Clearn GE, Vandenberg SG, Johnson RC, Rashad MN. Cognitive abilities: Use of family data as a control to assess sex and age differences in two ethnic groups. International Journal of Aging and Human Development 1975;6(3):261–76. doi:10.2190/BBJP-XKUG-C6EW-KYB7.

35. Rouleau I, Salmon DP, Butters N, Kennedy C, McGuire K. Quantitative and qualitative analyses of clock drawings in Alzheimer’s and Huntington’s disease. Brain and Cognition 1992;18(1):70–87. doi:10.1016/0278-2626(92)90112-Y.

36. Gorgolewski KJ, Varoquaux G, Rivera G, Schwarz Y, Ghosh SS, Maumet C, Sochat VV, Nichols TE, Poldrack RA, Poline JB, Yarkoni T, Margulies DS. NeuroVault.org: a web-based repository for collecting and sharing unthresholded statistical maps of the human brain. Frontiers in Neuroinformatics 2015;9(8):1–9. doi:10.3389/fninf.2015.00008.

37. Tosun D, Landau S, Aisen PS, Petersen RC, Mintun M, Jagust W, Weiner MW, Alzheimer’s Disease Neuroimaging Initiative. Association between tau deposition and antecedent amyloid-*β* accumulation rates in normal and early symptomatic individuals. Brain 2017;140(5):1499–512. doi:10.1093/brain/awx046.

38. Jack CR, Wiste HJ, Schwarz CG, Lowe VJ, Senjem ML, Vemuri P, Weigand SD, Therneau TM, Knopman DS, Gunter JL, Jones DT, Graff-Radford J, Kantarci K, Roberts RO, Mielke MM, Machulda MM, Petersen RC. Longitudinal tau PET in ageing and Alzheimer’s disease. Brain 2018;141(5):1517–28. doi:10.1093/brain/awy059.

39. Harrison TM, La Joie R, Maass A, Baker SL, Swinnerton K, Fenton L, Mellinger TJ, Edwards L, Pham J, Miller BL, Rabinovici GD, Jagust WJ. Longitudinal tau accumulation and atrophy in aging and Alzheimer disease. Annals of Neurology 2019;85(2):229–40. doi:10.1002/ana.25406.

40. Buckley RF, Mormino EC, Rabin JS, Hohman TJ, Landau S, Hanseeuw BJ, Jacobs HIL, Papp KV, Amariglio RE, Properzi MJ, Schultz AP, Kirn D, Scott MR, Hedden T, Farrell M, Price J, Chhatwal J, Rentz DM, Villemagne VL, Johnson KA, Sperling RA. Sex differences in the association of global amyloid and regional tau deposition measured by positron emission tomography in clinically normal older adults. JAMA Neurology 2019;doi:10.1001/jamaneurol.2018.4693.

41. Barnes LL, Wilson RS, Bienias JL, Schneider JA, Evans DA, Bennett DA. Sex differences in the clinical manifestations of Alzheimer disease pathology. Archives of General Psychiatry 2005;62(6):685–91.

42. Howell JC, Watts KD, Parker MW, Wu J, Kollhoff A, Wingo TS, Dorbin CD, Qiu D, Hu WT. Race modifies the relationship between cognition and Alzheimer’s disease cerebrospinal fluid biomarkers. Alzheimer’s Research & Therapy 2017;9(88):1–10. doi:10.1186/s13195-017-0315-1.

43. Graff-Radford NR, Besser LM, Crook JE, Kukull WA, Dickson DW. Neuropathologic differences by race from the National Alzheimer’s Coordinating Center. Alzheimer’s & Dementia 2016;12(6):669–77. doi:10.1016/j.jalz.2016.03.004.

44. Lee CM, Jacobs HIL, Marquié M, Becker JA, Andrea NV, Jin DS, Schultz AP, Frosch MP, Gómez-Isla T, Sperling RA, Johnson KA. ^18^F-Flortaucipir binding in choroid plexus: related to race and hippocampus signal. Journal of Alzheimer’s Disease 2018;62(4):1691–702. doi:10.3233/JAD-170840.

45. Bilgel M, An Y, Helphrey J, Elkins W, Gomez G, Wong DF, Davatzikos C, Ferrucci L, Resnick SM. Effects of amyloid pathology and neurodegeneration on cognitive change in cognitively normal adults. Brain 2018;8(August):2475–85. doi:10.1093/brain/awy150.

46. Ossenkoppele R, Schonhaut DR, Scholl M, Lockhart SN, Ayakta N, Baker SL, O’Neil JP, Janabi M, Lazaris A, Cantwell A, Vogel J, Santos M, Miller ZA, Bettcher BM, Vossel KA, Kramer JH, Gorno-Tempini ML, Miller BL, Jagust WJ, Rabinovici GD. Tau PET patterns mirror clinical and neuroanatomical variability in Alzheimer’s disease. Brain 2016;139(5):1551–67. doi:10.1093/brain/aww027.

47. Saint-Aubert L, Lemoine L, Chiotis K, Leuzy A, Rodriguez-Vieitez E, Nordberg A. Tau PET imaging: present and future directions. Molecular Neurodegeneration 2017;12(19):1–21. doi:10.1186/s13024-017-0162-3.

48. Ashburner J, Friston KJ. Spatial transformation of images. In: Frackowiak RSJ, Friston KJ, Frith C, Dolan R, Mazziotta JC, eds. Human Brain Function. Academic Press USA; 1997:43–58.

49. Jenkinson M, Bannister P, Brady M, Smith S. Improved optimization for the robust and accurate linear registration and motion correction of brain images. NeuroImage 2002;17(2):825–41. doi:10.1006/nimg.2002.1132.

50. Jenkinson M, Beckmann CF, Behrens TE, Woolrich MW, Smith SM. FSL. NeuroImage 2012;62(2):782–90. doi:10.1016/j.neuroimage.2011.09.015.

51. Zhou Y, Resnick SM, Ye W, Fan H, Holt DP, Klunk WE, Mathis CA, Dannals R, Wong DF. Using a reference tissue model with spatial constraint to quantify [^11^C]Pittsburgh compound B PET for early diagnosis of Alzheimer’s disease. NeuroImage 2007;36(2):298–312. doi:10.1016/j.neuroimage.2007.03.004.

52. Thomas BA, Cuplov V, Bousse A, Mendes A, Thielemans K, Hutton BF, Erlandsson K. PETPVC: a toolbox for performing partial volume correction techniques in positron emission tomography. Physics in Medicine and Biology 2016;61(22):7975–93. doi:10.1088/0031-9155/61/22/7975.

53. Avants BB, Tustison NJ, Song G, Cook Pa, Klein A, Gee JC. A reproducible evaluation of ANTs similarity metric performance in brain image registration. NeuroImage 2011;54(3):2033–44. doi:10.1016/j.neuroimage.2010.09.025.

54. Gorgolewski K, Burns CD, Madison C, Clark D, Halchenko YO, Waskom ML, Ghosh SS. Nipype: a flexible, lightweight and extensible neuroimaging data processing framework in python. Frontiers in Neuroinformatics 2011;5(13):1–15. doi:10.3389/fninf.2011.00013.

55. Allen EA, Erhardt EB, Calhoun VD. Data visualization in the neurosciences: overcoming the curse of dimensionality. Neuron 2012;74(4):603–8. doi:10.1016/j.neuron.2012.05.001.

56. Pinheiro JC, Bates DM, DebRoy S, Sarkar D, R Core Team. nlme: Linear and nonlinear mixed effects models. 2015. URL: http://cran.r-project.org/package=nlme.

57. Xie Y. Dynamic documents with R and knitr. 2nd ed.; Chapman and Hall/CRC; 2015. ISBN 978-1498716963.

58. Xie Y. knitr: A general-purpose package for dynamic report generation in R. 2018.

59. Zhu H. kableExtra: Construct complex table with ’kable’ and pipe syntax. 2019.

60. Hlavac M. stargazer: Well-formatted regression and summary statistics tables. 2018. URL: https://cran.r-project.org/package=stargazer.

61. Wickham H. ggplot2: Elegant Graphics for Data Analysis. Springer-Verlag New York; 2016. ISBN 978-3-319-24277-4.

62. Kassambara A. ggpubr: ‘ggplot2’ based publication ready plots. 2018. URL: https://cran.r-project.org/package=ggpubr.

63. Kurtzer GM, Sochat V, Bauer MW. Singularity: scientific containers for mobility of compute. PloS ONE 2017;12(5):e0177459. doi:10.1371/journal.pone.0177459.

64. Notter M, Gale D, Herholz P, Markello R, Notter-Bielser ML, Whitaker K. AtlasReader: A Python package to generate coordinate tables, region labels, and informative figures from statistical MRI images. Journal of Open Source Software 2019;4(34):1257. doi:10.21105/joss.01257.

65. Talairach J, Tournoux P. Co-planar stereotaxic atlas of the human brain. New York, NY: Thieme; 1988.

66. Lacadie CM, Fulbright RK, Rajeevan N, Constable RT, Papademetris X. More accurate Talairach coordinates for neuroimaging using non-linear registration. NeuroImage 2008;42(2):717–25. doi:10.1016/j.neuroimage.2008.04.240.

